# SOL3D: Soft-lithography on 3D vat polymerised moulds for fast, versatile, and accessible high-resolution fabrication of customised multiscale cell culture devices with complex designs

**DOI:** 10.1101/2022.02.22.481424

**Authors:** Cathleen Hagemann, Matthew C. D. Bailey, Eugenia Carraro, Valentina Maria Lionello, Noreen Khokhar, Pacharaporn Suklai, Carmen Moreno-Gonzalez, Kelly O’Toole, George Konstantinou, Sudeep Joshi, Eleonora Giagnorio, Mads Bergholt, Albane Imbert, Francesco Saverio Tedesco, Andrea Serio

**Affiliations:** Centre for Craniofacial & Regenerative Biology, King’s College London, London SE1 9RT, UK; The Francis Crick Institute, London NW1 1AT, UK; Department of Cell and Developmental Biology, University College London, London WC1E6DE, UK; Dubowitz Neuromuscular Centre, UCL Great Ormond Street Institute of Child Health & Great Ormond Street Hospital for Children, London, UK; Neurology IV - Neuroimmunology and Neuromuscular Diseases Unit, Fondazione IRCCS Istituto Neurologico Carlo Besta, Milan 20133, Italy; Randall Centre for Cell and Molecular Biophysics, King’s College London, London SE1 1UL, UK; Dementia Research Institute (UK DRI)

## Abstract

Cell culture devices, such as microwells and microfluidic chips, are designed to increase the complexity of cell-based models whilst retaining control over culture conditions and have become indispensable platforms for biological systems modelling. From microtopography, microwells, plating devices and microfluidic systems to larger constructs for specific applications such as live imaging chamber slides, a wide variety of culture devices with different geometries have become indispensable in biology laboratories. However, while their application in biological projects is increasing exponentially, due to a combination of the techniques and tools required for their manufacture, and the physical science background sometimes needed, the design and fabrication of such devices directly by biological labs remains a relatively high investment in terms of costs, use of facilities, needed collaborations and time. Whilst commercially available systems are available, these are also often costly, and importantly lack the potential for customisation by each single lab. This combination of factors still limits widespread application of microfabricated custom devices in most biological wet labs.

Capitalising on recent important advancements in the fields of bioengineering and microfabrication, and taking advantage of low-cost, high-resolution desktop resin 3D printers combined with PDMS soft lithography, we have developed an optimised low-cost and highly reproducible microfabrication pipeline, capable of generating a wide variety of customisable devices for cell culture and tissue engineering in an easy, fast reproducible way for a fraction of the cost of conventional microfabrication or commercial alternatives. This protocol is designed specifically to be a resource for biological labs with little to none prior exposure to these fields technique and enables the manufacture of complex devices across the µm to cm scale.

We provide a ready-to-go pipeline for the efficient treatment of resin-based 3D printed constructs for PDMS curing, using a combination of curing steps, washes and surface treatments. Together with the extensive characterisation of the fabrication pipeline, we show the utilization of this system to a variety of applications and use cases relevant to biological experiments, ranging from micro topographies for cell alignments to complex multi-part hydrogel culturing systems.

This methodology can be easily adopted by any wet lab, irrespective of prior expertise or resource availability and will enable the wide adoption of tailored microfabricated devices across many fields of biology.

## 1. Introduction

Stem cell-based models are an invaluable resource, which allows the study of nearly any cell type *in vitro*^1–4^. The advent of cellular reprogramming and subsequent access to patient-derived stem cell models have also galvanised their position as an ideal tool to investigate cellular processes in health and disease^5–9^. While stem cell models offer significant control over the identity of cultured cell types, the conventional culture systems used for them typically lack the ability to control key parameters of the culture itself, which greatly influence the analyzed biological processes. These parameters include the relative position of the cultured cells, grouping, cell-cell and cell-material interactions. Although some commercially available devices allow for some control over culture conditions, they are often non-customizable and limited in their application to different research questions. Therefore, it is desirable for biologists to design and manufacture their own custom cell culture devices.

Several bioengineering strategies have been developed to create custom-engineered culture environments that direct the cell’s environment ^10,11^ . To achieve this, two main requirements are necessary: a suitable cell culture material and a method for shaping it into the desired form. PDMS (polydimethylsiloxane) is one of the most used materials for cell culture devices due to its numerous advantages. These include optical clarity, making it suitable for microscopy, and tunable stiffness ranging from 800 kPa to 10 MPa. PDMS also allows for a wide range of chemical modifications. However, PDMS is a porous material that absorbs small molecules and growth factors, necessitating single usage of PDMS constructs and frequent preparation of new devices. For this reason, fast and reliable manufacturing of these devices is often a crucial limit step in several experimental pipelines.

Most microdevices used in cell culture experiments require features ranging from several millimeters to micrometers, depending on the size of the cells, the required volume of the culture medium and needed experimental paradigm.

Several microfabrication techniques have been adapted over the years to the needs of the biology research community, and as a result the use of micropatterned substrates, microchamber devices and other engineered substrates has increased exponentially. One of the most versatile combination of techniques to obtain PDMS devices for biological experiments is photolithography coupled with soft-lithography^12,13^ . Briefly, photolithography is based on the deposition of layers of UV-sensitive photoresist of desired thickness, which is then exposed to a UV source with either a photomask or by direct laser writing to create the desired design, before developing the exposed photoresist. This process can be repeated for multiple layers and allows to create 2.5D designs (i.e. different planar structure of different thickness stacked to form one single set of features)

While they can be used to create advanced *in vitro* culture systems with micron-scaled features, photolithography based pipelines also present some limitations; for example, they can only create a single layer at a time with a given height determined by the photoresist layer’s properties, and are generally limited in the aspect ratio of the features that they can create. As a result, features are limited to 2.5D designs with defined thickness, lacking 3D volumes, curves, or interconnected shapes (Diagram 1B). Moreover, generating multi-layer constructs for complex features involves multiple photolithographic steps, which can be time-consuming, prone to errors, and costly (Supplement Diagram 1 A-B).

**Diagram 1:**
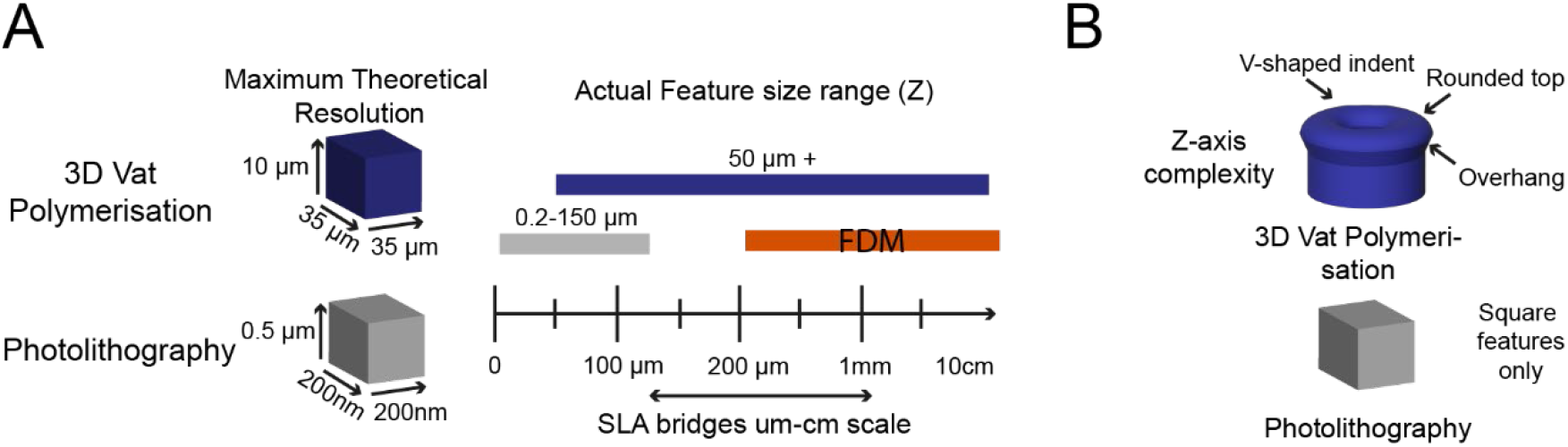
Comparison between feature constructing techniques; photolithography and UV resin vat polymerisation. **(A)** Scale comparison of 3D printing methods UV resin vat polymerisation, FDM and photolithography. Comparison of maximal resolution achieved with UV resin vat polymerisation, with current printers, and photolithography. (B) Representation of feature designs achievable with UV resin vat polymerisation and photolithography. UV resin vat polymerisation offers a wide range of feature dimensions and complexity compared to photolithography which underlies technical limitations for designs.

While these limitations can be obviated by recent improvements and optimisations in photolithography, such as grey scale photoresist^14,15^ and high resolution 2-photon based lithograph^16^, the result is often increased complexity in the fabrication process, and in costs or availability of the necessary instruments.

Consequently, it remains challenging for wet labs to perform rapid prototyping of user-handleable macroscale devices with microscale features for cell culture applications. Additionally, specialized facilities and expertise are often required for these techniques, limiting access for many biology focused groups and potentially creating a bottleneck for wider adoption of microfabrication. (e.g., imaging chambers for microscopy with fixed geometry or volume). 3D printing technology has emerged as an accessible and adaptable tool for fast prototyping and fabrication of small objects. Alongside their increasing availability, rapid technological advancements in 3D printers have led to the development of several open-source projects aiming to enable any wet lab to create and adopt critical and innovative modeling strategies ^17–22^. This is particularly important when considering the challenges experienced by laboratories in less developed countries in sourcing equipment or specific consumables, or the sometimes-steep practical barrier that some labs encounter when venturing into cell culture and biology from a different field.

There are a variety of different 3D printing techniques, ranging from filament deposition to vat polymerisation of resins. Most of these basic techniques are commercially available but more complex and intricate techniques have been developed over time, for example two-photon based microfabrication^23, 24^. Although these custom-built, high-resolution setups offer significant advancements, our focus here will be on vat polymerization as it represents the most accessible and cost-effective form of 3D printing with a µm-scale resolution.

UV resin vat polymerisation is a specific type of 3D printing based on light-curable resin, that with recent advancements in maximal resolution and pricing, shows the potential to bridge the gap between μm-resolution photolithography and mm-resolution fused deposition modelling (FDM) 3D printers **(Diagram 1A, Supplementary 17)**, whilst remaining economically accessible to any lab **(Figure S1).** Because of the positive characteristics UV vat polymerisation provide while remaining accessible, coupled with the rapid and easy fabrication of complex shapes in 3D, UV resin vat polymerisation represents in theory an ideal technique to create bespoke culture vessels, inserts and other devices to increase complexity within biological experiments without sacrificing control over culture conditions ^25, 26^.

Unfortunately, most commercially available resins for UV resin vat polymerisation are cytotoxic and cannot be used for cell culture applications^27^. Additionally, the composition of these resins is often proprietary, and conversion or production of biocompatible resins requires skills limited to dedicated chemistry laboratories^28^. Some biocompatible resins are commercially available; however, they tend to be sold at a much higher cost than even high-resolution resin, and more than the actual printers in some cases (e.g Phrozen sonic mini 4K printer = GBP 365 ^29^, 1L Zortrax Raydent Crown and Bridge resin = GBP 392 ^30^), undermining the applicability of 3D SLA printing for cell culture purposes **(Figure S1)**, even though these prices can be variable and might change over time but are here used to exemplify the lower cost. Moreover, unlike PDMS and other silicon-based materials used for soft-lithography resins do not have tuneable stiffness and generally are not optically clear and cannot be used for microscopy studies.

One possible solution to these problems would be to combine PDMS soft lithography with UV resin vat polymerised 3D printed moulds, and effectively employ 3D printing in lieu of photolithography in conventional pipelines. However, curing of PDMS on vat polymerisation resin prints can be quite challenging as acrylates and triorganophosphate photo-initiators, constituents of most commercially available resins, inhibit PDMS polymerisation^31–34^. Furthermore, vat polymerisation -resin-induced PDMS curing inhibition makes demoulding difficult and can result in leaching of cytotoxic uncured PDMS monomers into the cell culture medium of even successfully demoulded designs^35^, making the devices unusable for cell culture applications.

To overcome these challenges and facilitate the production of complex 3D constructs suitable for cell culture, several successful post-processing and coating approaches have been attempted^36–38^. However, these protocols generally involve either long heat and detergent treatments^39^, which often cause print deformation, or expensive techniques^40^, not accessible to every lab. One example of the latter, coating of UV resin vat polymerised prints with parylene is effective in creating usable moulds, and is sufficient to overcome curing inhibition of PDMS^41^, but require the use of specialised equipment and adds another potentially technically challenging step to optimise.

Driven initially by our own experience with adapting microfabrication techniques to hard biological questions, and inspired by the many recent technical advancements by different groups, we aimed to create an optimized and universally effective pipelines that would include the production and post-processing protocol for **so**ft **l**ithography on **3D** vat polymerised moulds (**SOL3D**), using a low-cost commercially available printer and materials.

To demonstrate the applicability of this method to several different biological experiments and provide an effective blue-prints for other labs that don’t have expertise in microfabrication, we demonstrated its use to develop customisable culture devices ranging from µm to mm and cm scale, with complex 3D shapes or and micro topographies. Together with the detailed protocols, we also provide the designs for each device showcased, which can be customised to fit different experimental needs.

## 2. Results

### 2.1 Optimisation of PDMS curing on 3D SLA printed moulds

To overcome the current barriers preventing the integration of 3D vat polymerisation for the fabrication of tissue culture constructs in biology labs, we aimed to optimise an easy-to-implement and widely applicable protocol to enable efficient PDMS curing on vat polymerised moulds using commercially available equipment. We, therefore, tested a variety of commercially available resins **(Table 3)** subsampling different manufacturers together with a commercially available high-resolution 3D vat printer (300-400 GBP retail price, Phrozen 4K Sonic Mini or Anycubic Photon S equivalent to approx. 2 batches of a monoclonal antibody for immunostaining).

First, we verified the previously reported cytotoxicity of each resin following conventional post-processing steps (isopropanol washing, UV curing), either in an untreated state or with supplementary heat treatment, washing and UV sterilisation, by co-culturing chips of resin with induced pluripotent stem cells (iPSC)-derived motor neurons (MNs) **(Figure S2)**. Additionally, we tested a grad 2a biocompatible resin used for dental implants. After 4 days of coculture with this resin, the iPSC-derived MNs show despite different washing protocols toxic effects, and a clear visual increase in debris compared to the control well (Figure S16). As none of the resins were suitable for cell culture applications in direct use, we then focused on optimising the post-print processing protocol for resin moulds testing different parameters across three main steps: resin washing, print coating, and PDMS heat treatment curing **(Figure 1A)**, to find an easy and fast method overcoming PDMS curing inhibition, as no standardised post-processing protocol exists **(Figure 1B)**. Resins were printed using modified manufacturer’s settings for the recommended printer (either Anycubic or Phrozen) **(Table 3) (Figure 1C).** We tested PDMS curing on the moulds at 6 different time points (2h, 4h, 6h, 18h, 22h, 24h) and considered the sample conditions not optimal for curing if the process took over 30h.

**Figure 1:**
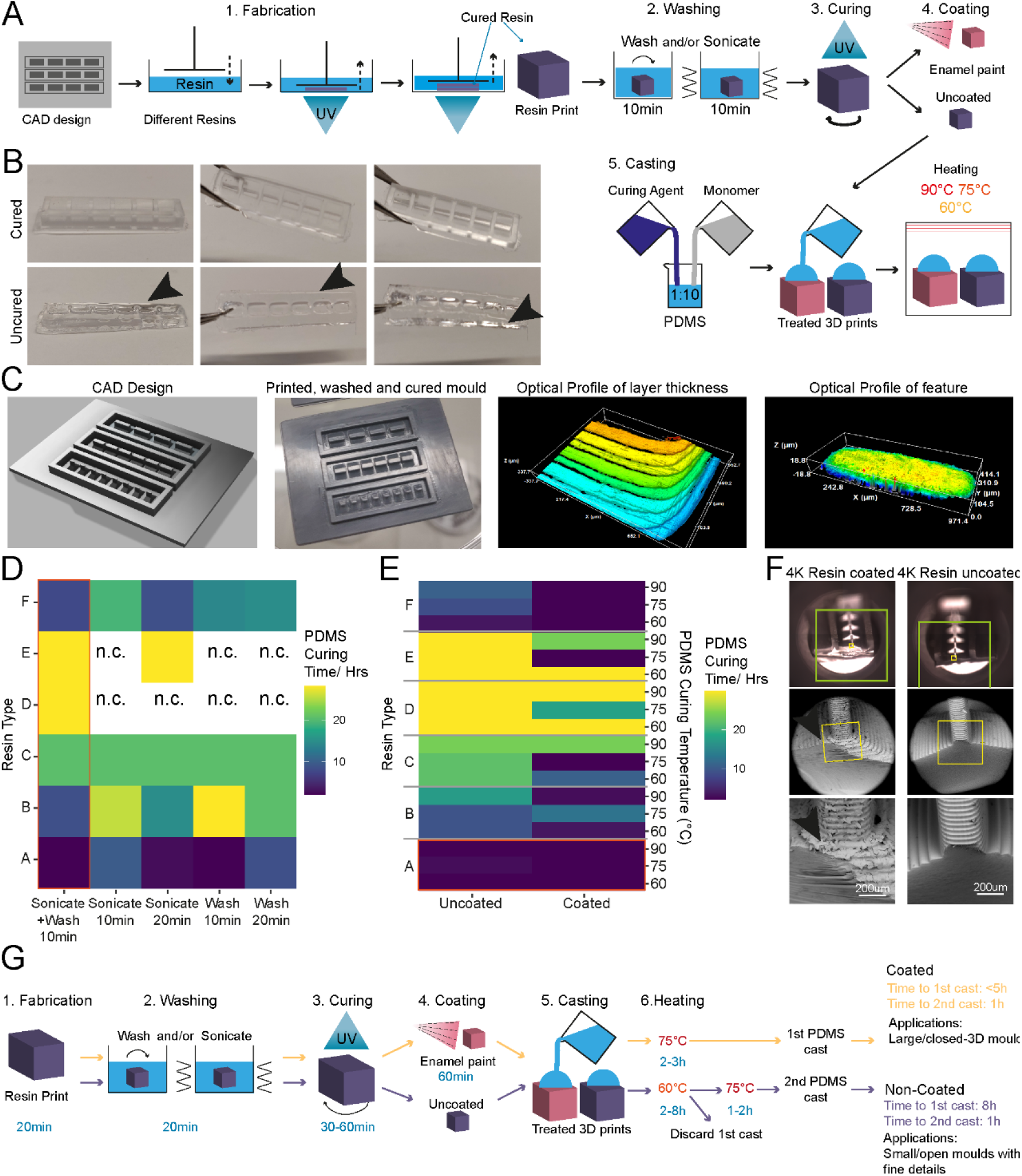
Enamel paint coating facilitates rapid PDMS curing on 3D printed moulds. (A) Schematic overview of the investigation strategy to establish a protocol for PDMS curing on 3D printed moulds (B) Representative images of PDMS casts removed from printed devices classified as cured or uncured. Arrows highlight liquid PDMS. (C) Representative images of a CAD of 3D printed moulds, the completed print, and surface optical profiles of layer thickness and feature dimensions. (D) Heatmap of PDMS curing time by resin type for different washing conditions. (E) Heatmap of PDMS curing time by resin type for different PDMS curing temperatures (right y-axis) and different SLA print coatings. (F) Representative SEM images of uncoated (right) and enamel paint coated prints (left), arrows highlight the paint layer. (G) Schematic overview of optimised fabrication, post processing and PDMS casting protocols with (yellow) and without (purple) enamel coating.

The isopropanol washing step is designed to remove excess uncured resin from the printed moulds. We tested two different methods for removal, sonication and stirring, alone or in combination, each for 10 minutes, and found that post-printing washing conditions in isolation have a modest effect on PDMS curing time, but a combinatorial treatment was beneficial. Therefore, all subsequent experiments were performed using sequential treatment with sonication and washing (10 minutes each). Interestingly, we found that resin selection had more impact on curing time than washing itself, with resins A and F performing the best **(Figure 1D)**. Washing of resin E and D was unsuccessful in most conditions, due to the amount of uncured resin adhering to the print from improper printing, and subsequent analysis of PDMS curing on these samples would bias the curing time if curing can take place at all.

It has been suggested that curing inhibition on resin can result from vaporised acrylate monomers ^36^, which are components of most resins, released into the PDMS during heating. We reasoned that either blocking the contact sites between acrylates and PDMS or reducing the release of acrylates from the resin during curing could be sufficient to allow efficient curing of PDMS on the moulds. To test these hypotheses, we used commercially available enamel paint to homogenously coat the washed 3D prints with an airbrushing system, forming a protective barrier between the PDMS and resin. We then compared the PDMS curing time of coated prints to uncoated prints at 3 different temperatures (60, 75 and 90°C), which allowed us to identify the role of temperature on acrylate release and PDMS curing. These experiments showed that enamel paint coating enabled PDMS curing not only on the surface but throughout the whole cast, and decreased PDMS curing times for all resins. It is important to note that Resin A is a special case, as it showed good PDMS curing performance with and without coating **(Figure 1E)**, permitting the use of our post-processing protocol with and without enamel paint. These two protocols differ not only in the coating but also in the curing temperature used, which impacts the overall manufacturing times. The benefit of the missing paint layer of the non-coating protocol is that it allows the manufacture of detailed PDMS moulds (features <300 µm) **(Figure 1G)**. Overall, lower temperatures improved PDMS curing times on resins and the variation in the effect of temperature on coated samples was negligible. Additionally, we observed that high temperatures (90°C) resulted in a significant warping of the print. These unwanted effects were less prominent at lower temperatures (60-75 °C), which were still effective enough to cure PDMS.

It has also been reported that acrylate monomers and photo-initiators, which are resin components, can leach from the resins into PDMS, impacting the biocompatibility of cast constructs ^35^. To verify that the cured PDMS samples did not have leachates of resins or enamel paint, we used Raman spectroscopy to characterise the chemical composition of samples from each post-processing condition and compared them to cured and uncured PDMS (**Figure S3**). This spectral analysis revealed no detectable carryover of resin constituents or paint into the casted PDMS and high similarity of resin casts to cured PDMS. To further validate these findings, we cocultured resin print cast PDMS substrates with iPSC-derived MNs. These co-cultures demonstrated good biocompatibility over longer culturing periods (**Figure S4**).

As introducing a layer of enamel coating on the mould devices could negatively impact fine feature sizes, we used SEM imaging on printed moulds with resin A to quantify whether print dimensions and surface roughness were significantly affected. Analysis showed that coated prints exhibited a thin layer of paint <100 µm and greater surface roughness compared to uncoated ones **(Figure S5)**. This limits the application of this method to features larger than 100 µm in any dimension **(Figure 1F)**.

Overall, we established a fast and robust post-processing pipeline, identified suitable resins and provide a ready-to-use protocol for 3D vat printing. We provide two modified versions of an effective post-processing protocol for **so**ft-**l**ithography on **3D** vat polymerised moulds, depending on the design and feature size **(Figure 1G)**.

### 2.2 SOL3D fabrication allows the generation of complex 3D-shaped stencils for precise control of cell positioning and grouping within open wells

#### 3D printed stencil-aided dry plating devices to control cell location and number within standard well plates

Conventional open well culture systems generally do not allow control over cell position, grouping, and numbers in an easy and reproducible fashion, limiting the complexity of *in vitro* modelling experiments. Several techniques are available to overcome these limitations and to create precise arrangements of cells within culture vessels -from microfluidic devices to cell bioprinting-, however, most rely on creating compartmentalised structures which limit the manipulation of the cells granted by the open well systems.

A different approach that allows both to increase complexity within conventional culture vessels and to maintain an open well system are stencil-like plating devices ^42, 43^. These systems are temporary structures that guide the organisation of cells within an open well, although at present they suffer from the same fabrication limitations as the above-mentioned strategies. Moreover, this method is especially affected by the technical limitations of feature sizes and aspect ratios dictated by photolithography, resulting in thin devices that are difficult to handle and have limited customisation possibilities **(Figure S6)**.

We decided to use these types of devices, using human induced pluripotent stem cell-derived motor neurons (MNs) as a cell model system, for an initial proof-of-principle of our optimised SOL3D protocol, based on an engineered platform we recently developed for MN cultures using a micropatterned substrate to facilitate axonal elongation ^44^.

We combined and optimised this platform with our SOL3D moulding protocol to create a tailored plating strategy for investigating hiPSC-derived MN behaviour with control over cell location and orientation. We designed moulds for casting PDMS stencil-well devices, rectangular extruded features with funnel-shaped media reservoirs as complex 3D features to ease cell seeding. This optimised design permits rapid and facile manual seeding as cells can settle into micro-sized wells in a suitable volume of medium to avoid excessive evaporation and cell death **(Figure 2 A-B)**.

**Figure 2:**
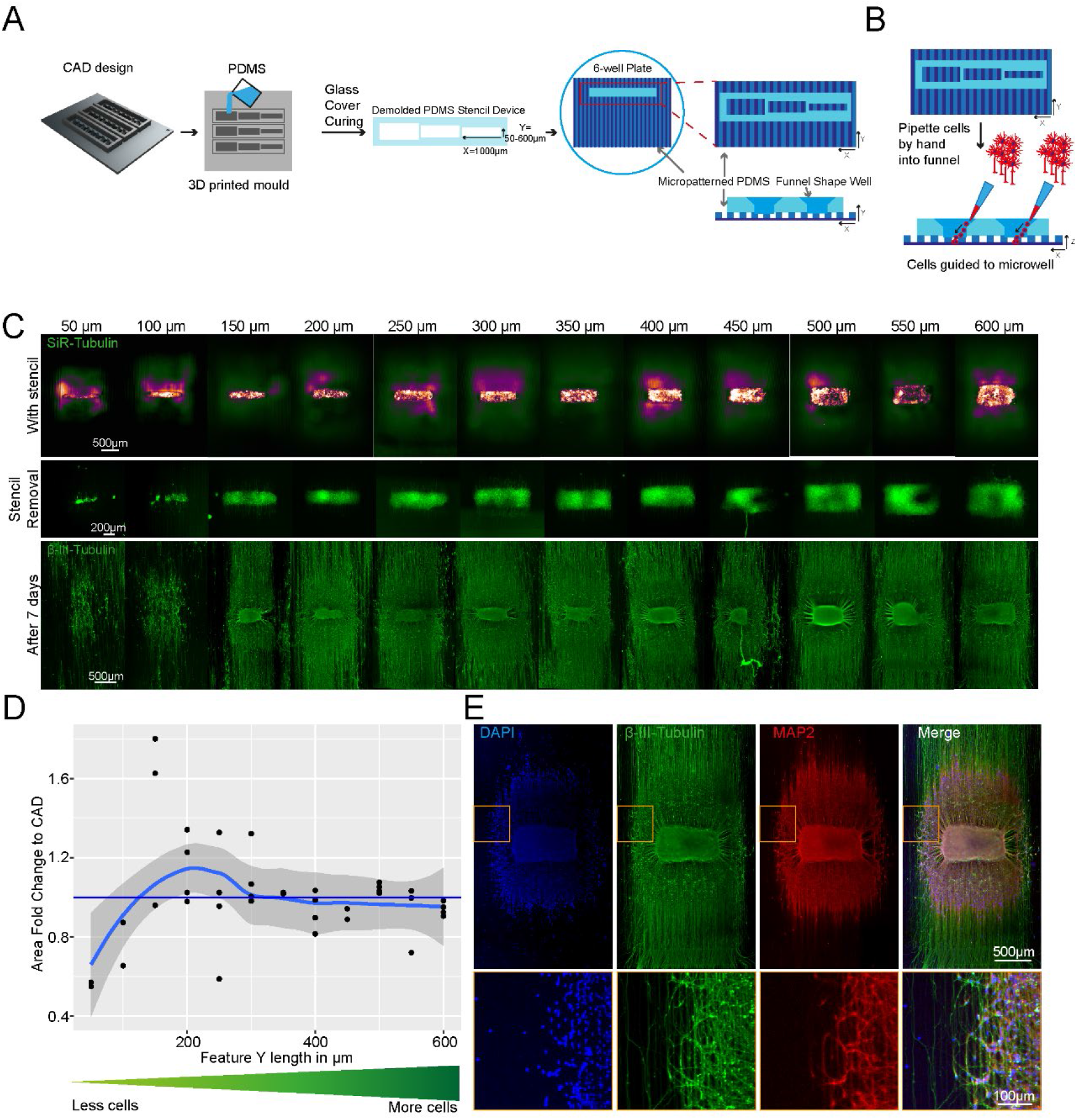
SLA 3D printing enables control over cell location and number in an open well. (A) Schematic overview of the investigation dry plating strategy to combine PDMS casts from 3D printed molds with PDMS microgroove substrate to control cell body location and number. Well sizes range from 600 µm x 1000 µm to 50 µm x 1000 µm in 50 µm intervals (B) Schematic overview of funnel shape well for easy manual seeding in microwells. (C) Representative images of stencil devices filled with pre stained (Silicon Rhodamine-tubulin) MN progenitors with device still in place (top), after device removal (middle), and axonal β-III Tubulin following fixation after 7 days of culture (bottom). (D) Comparison of seeded cell area after stencil removal to CAD specified values by fold change (E) Representative images of 3D aggregoid with 2D axon elongation stained with DAPI (1^st^ image), axonal β-III Tubulin (2^nd^ image), and Dendritic MAP-2 (3^rd^ image) simultaneously (4^th^ image).

PDMS stencils from these 3D moulds allow seeding by “dry plating”, whereby a stencil is placed in a dry conventional tissue culture plastic vessel and cells in suspension are manually pipetted in the stencil device, isolating the cell bodies from the residual well and allowing them to adhere at these specific positions. With the cell bodies secured, the stencil device can be removed and the whole well filled with culture medium, while the adhered cells remain in their specified position. For this “dry plating” process, a strong fluidic seal surrounding the PDMS stencil wells is necessary, requiring a flat surface between the stencil and substrate below. Without specific steps to adjust the surface roughness of prints, PDMS casts from 3D printed moulds are inherently rougher than those from micropatterned silicon wafers used for casting microfabricated PDMS devices **(Figure S7)**. We, therefore, implemented an additional clamping step before PDMS curing, using a silanised glass slide *(see M&M)* to cover the PDMS surface, which is in contact with air, taking advantage of the flat surface provided by the glass (**Figure S8A)**. We evaluated the efficiency of clamp-cured stencil fluid seals when placed on a PDMS micropatterned surface with 10 x 10 µm-grooves using a blue dye. An effective seal was achieved in all stencils cured using the additional clamping, denoted by dye reaching the microgroove substrate in the well area only and spreading within these specific grooves. Stencils cast without clamping showed uncontrolled dye spreading throughout the devices, verifying a lack of fluid seal **(Figure S8B).**

We were then able to use the optimised stencil devices to answer a biological question and investigate the minimum number of iPSC-MNs required to form a self-organised 3D neural aggregate on microgrooves for axonal elongation, a process determined by chemotaxis and topography. To achieve this, we used the above-described stencils with a funnel shaped reservoir and rectangular wells, varying in Y-dimension to reduce the stencil well size and control cell amount. The well dimensions were homogenous and faithful to CAD specifications throughout the print sizes down to 50 µm in Y **(Figure S9)**. These PDMS stencil-well devices were placed on the extra cellular matrix (ECM) coated and dried micropatterned surface with axonal guidance grooves ^44^, and the iPSC-derived MN cell suspension was manually pipetted into the dry wells of the device. To avoid potential air pockets in the smaller wells, as it is common for non-functionalised PDMS, we performed oxygen plasma treatment on stencils prior to cell “dry plating” (**Figure S10).** Compact rectangular “aggregoids” (i.e. 3D cell clusters generated by reaggregating single cells from a culture) with decreasing size were achieved during seeding and were maintained following device removal. Staining with β-III-tubulin after 7 days in differentiation medium revealed that wells with a size of down to 150 µm provide suitable cell numbers for aggregate formation. However, the two smallest well sizes did not provide the environment for aggregate formation and cells migrated across the topography **(Figure 2C-D).** Subsequent staining with compartment specific markers showed a clear separation between dendrites and axons in the open well devices of compact aggregoids **(Figure 2E)**.

In summary, stencil-well devices cast with SOL3D can be used to control cell location in an open well, facilitate control over different cell numbers in the same device, and enable cell compartment specific investigations.

#### Spatial, temporal and morphological control over cell-cell interaction using tailored plating devices

We next explored the potential to use SOL3D for more complex cultures, incorporating different cell types and plating timepoints.

First, to plate multiple cell populations within the same devices we sought to take advantage of PDMS natural hydrophobicity coupled with the large rectangular well design and funnel shaped well profile showed in **Figure 2A**. The high contact angle between media and hydrophobic PDMS allows to achieve confinement of the different cell suspension droplets, which generates enabling complete fluidic separation between adjacent wells containing different cell populations, even with manual plating. We sought to utilise this strategy to simultaneously plate different iPSC-derived MNs populations within the same tissue culture well in different spatially separate pockets of the PDMS device. For this we used fluorescent RFP^+^ and untransfected MNs, which we plated manually within adjacent pockets in the same device placed on an ECM coated well of a 6-well plate. The different cell populations were left to adhere for 2h before the plating device was removed. After removing the device and further cell culture for 72 hours, all MNs were stained with a silicon rhodamine tubulin dye (here depicted in green for visualisation), to visualise all neurites and cell bodies. A line graph analysis across the whole device showed that all wells contained MNs (RFP^+^ and RFP^-^/ Tubulin^+^) and in every second well, RFP^+^ cells were present **(Figure 3A-B)**, demonstrating multi cell type seeding in confined predetermined spatial groups within a single device.

**Figure 3:**
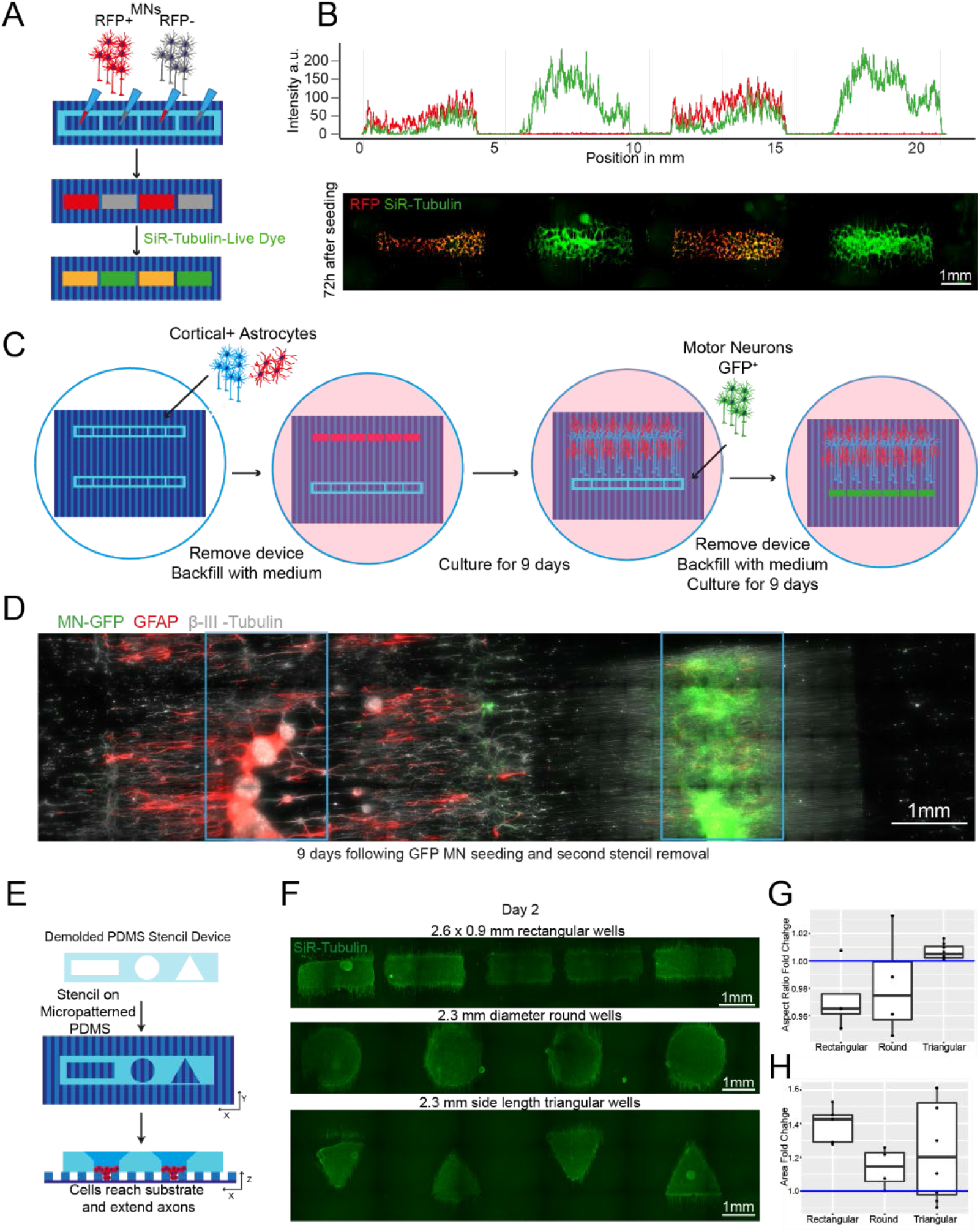
Plating devices enable spatiotemporal control of cell plating with different geometries for construction of complex neural circuits. (A) Schematic overview of alternate seeding of RFP and non-RFP+ motor neurons in the same device and the following live cell staining (B) Representative line profile of stained across the well showing segregation of individual populations to their designated wells – RFP+ only in wells 1 and 3, but SiR-tubulin+ (here in green) in all wells (top). Representative fluorescence images of stained RFP+/- motor neurons (bottom) (C) Schematic overview of the multidevice protocol for constructing a neural circuit using 2 stencil devices and 3 different cell types (MNs, cortical and astrocytes) with seeding performed at different time points (D) Composite of the complete circuit after 19 days of culture. GFP transfected motor neurons = green, Glial Fibrillary Acidic Protein (GFAP) identifies astrocytes, β-III – Tubulin identifies cortical neurons and the tubulin in GFP+ motor neurons. Blue device well shapes overlaid for illustrative purposes. (E) Schematic overview of protocol for manipulating aggregate geometry in combination with existing microgroove (F) Representative images of SiR-tubulin live-cell stained motor neuron aggregoids at day 2. (G) Boxplot of aggregoid aspect ratio fold change by shape between CAD (blue line) and day 2 of culture from β-III-Tubulin channel (top). (H) Boxplot of aggregoid area fold change by shape between CAD (blue line) and day 2 of culture from β-III Tubulin channel.

We then sought to further increase the complexity of our *in vitro* cultures by seeding multiple cell types at different time points within the same well, taking advantage of the efficient and reversible fluid seal between our devices and the culture plate. For this, we placed two rectangular plating devices, approximately 2mm apart from each other on an ECM-coated micropatterned surface within a well of a 6-well plate, as described above. Initially, one device was used to dry plate iPSC-derived cortical neurons^45^ and astrocytes^46^ in a 1:1 ratio and removed after 24h, while the other device was kept empty. The whole tissue culture well was then filled with differentiation medium and cultured for 9 days. During this time, cortical axons guided by the microtopography extended toward the empty device, which maintained its initial fluidic seal even surrounded by medium. On day 9, GFP^+^ MNs were seeded in the second device by first lowering the level of the medium within the well to be below the edge of the plating device, and then seeding the MNs in suspension directly within it. After allowing for cell attachment, the second device was also removed, the well refilled with fresh medium and cells cultured for further 9 days. The position of the different cell types was then verified using immunocytochemistry (ICC) for astrocytes (GFAP) and neurons (both MNs and cortical neurons, β-III-Tubulin), as cortical neurons could be identified by the overlap of GFP and β-III-Tubulin. Using these tailored removable SOL3D-generated plating devices we were able to easily plate 3 different cell types at 2 different time points within the same culture well, creating a complex neural circuit and demonstrating true spatiotemporal control over cell seeding in a cost-effective and highly adaptable fashion **(Figure 3C-D, Supplementary 11)**. Additionally, we demonstrated also multiple time point seedings within the same well using large format “nesting” plating devices which can be used to construct large-scale cell and tissue arrangements (**Figure S12).**

Next, we tested whether the ease of available prototyping using our optimised SOL3D protocol could enable investigation and manipulation of the fundamental behaviour of complex iPSC-derived MN cultures. It has been shown that aggregation of cells using different geometries has an influence on the signalling environment and patterning of aggregates^47^. We therefore designed and fabricated moulds for PDMS stencils with 3 different well shapes: rectangular, circular, and triangular, to create geometrically constrained neural aggregates. With the advantage of volumetric 3D printing, we were able to preserve the funnel reservoir and straight well design from previous moulds (**Figure 3E)**. Using these multi-shaped stencils, we seeded motor neuron progenitors (MNPs) as before on our micropatterned substrate and allowed axons to extend for 11 days (**Figure 3E)**. Here, MN aggregates maintained faithful area and aspect ratios to CAD specifications on day 2 after stencil removal. After 11 days, the groups also retained their specific geometry although showed slight changes in the aspect ratio and area over time **(Figure 3F-H).** Next, we used these PDMS stencils on nonpatterned and uncoated tissue culture plastics to avoid cell adherence, directing self-organisation of aggregate-like structures (**Figure S13 A)**. Here, we seeded cortical progenitors in Matrigel and were able to generate differently shaped aggregates demonstrated by SiR-tubulin live dye images after 24 hours **(Figure S13B).** SOL3D fabrication can therefore be used as a valid method of fabricating constructs for controlling cellular interactions in complex cultures of multiple geometries for both 2.5 (i.e. partially tridimensional adherent cultures) and 3D non-adherent cultures (e.g. aggregates) depending on the seeding substrate.

### 2.3 SOL3D fabrication allows the generation of micro topographies

In the initial experiments presented in **Figures 2 and 3**, we employed SOL3D-fabricated devices on top of microgrooves generated with conventional photolithography. As this method is not available to all labs, we then focused on achieving the same level of organisation within the culture but on a platform purely based on SOL3D-manufactured devices. While the 3D vat polymerisation printers do not have the same resolution as photolithographic equipment, we could not simply recreate the 10µm grooves.

Moreover, with current LCD-based illumination, repetitive patterns in close proximity to each other and close to the minimal resolution pose a challenge for 3D vat polymerisation printing, due to the illumination pattern and diffraction of the light.

However, the key parameter is the biological organisation rather than the material geometry, and we, therefore, focused on obtaining a design that can both be printed with SOL3D and achieves the same neurite orientation. We developed a design with different groove geometry and dimensions (**Figure 4A**), with a groove depth of 200µm and width of 100 µm. Optical profiling of the 3D prints reveals that the grooves are shallower than the CAD design, around 60um, and less wide, as expected (**Figure S13**). Despite these limitations, we tested if the topography is sufficient to align axonal elongation. We plated MN neural aggregates^44^ in a SOL3D-manufactured stencil device on SOL3D-manufactured grooves following the ‘dry plating’ protocol (**Figure 4B**). Visualisation of axons (β-III-Tubulin) after 7 days of culture revealed an alignment with the topography throughout the axonal length (**Figure 4C-D**).

**Figure 4:**
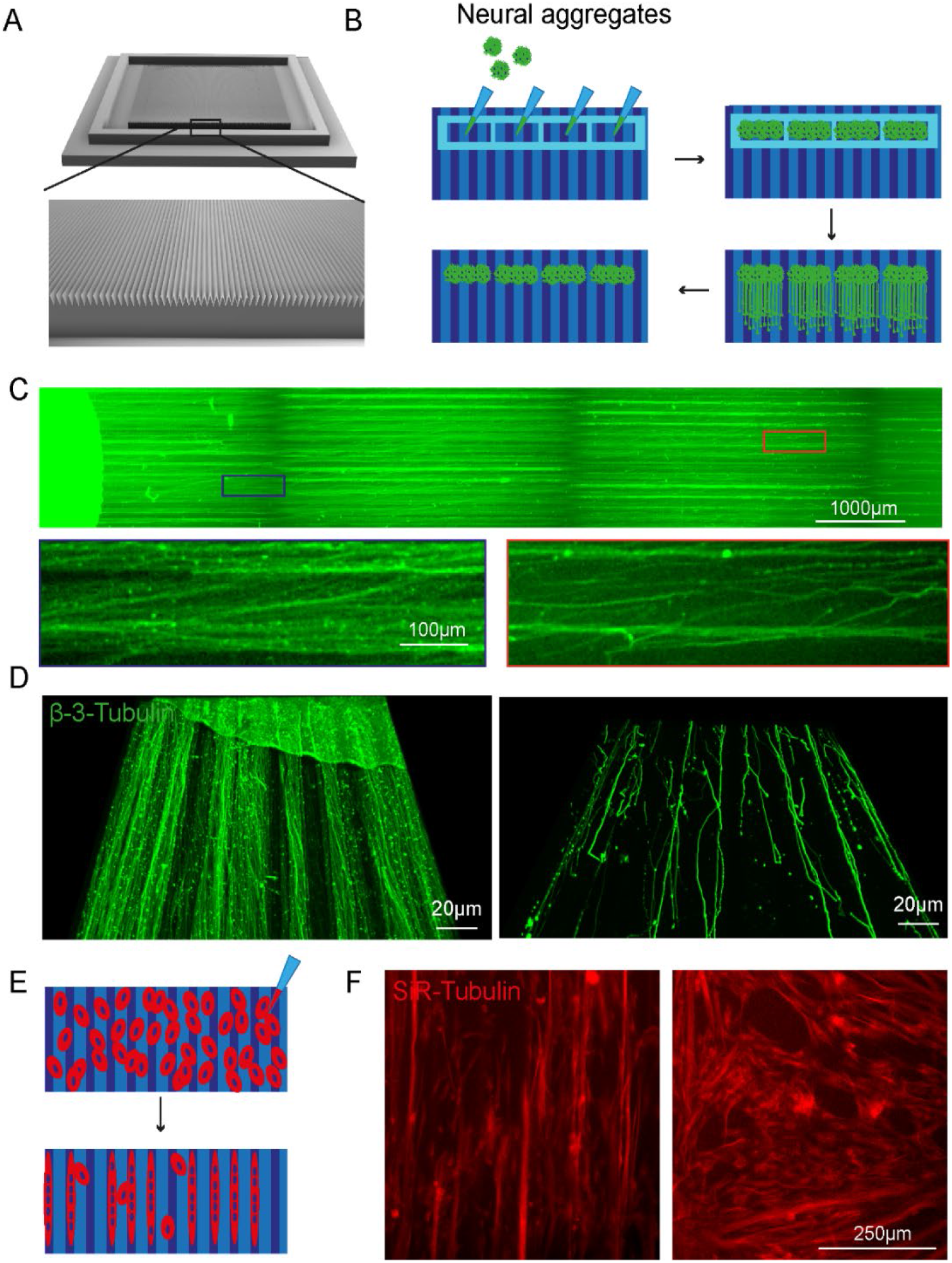
SOLID fabricated grooves guide axonal elongation and alignment of muscle fibres. (A) CAD design of triangular grooves 100um wide and 200um deep with 75° triangular spacing in between (B) Schematic overview of ‘dry’ neural aggregate seeding in PDMS stencil devices on the 3D printed grooves and axonal elongation. (C) Representative overview image of axons(β-III-Tubulin) on the SOLID fabricated grooves with magnifications of axons aligning with the topography in the proximal (blue) and distal (orange) compartment. (D) Representative 3D reconstruction of axons (β-III-Tubulin) on grooves in the proximal (left) and distal (right) compartment. Across compartment axonal alignment to the topography is given. (E)Schematic representation of myoblast seeding and differentiation on SOLID grooves. (F) At day 3 of differentiation myotubes (SiR-Tubulin) align to the given topography(left) compared to cells cultured on a non-patterned surface (right).

Another process benefitting from alignment to topographies is the formation of myotubes^48^. We seeded myoblasts on the SOL3D grooves and a flat control surface (**Figure 4E**). Cells were differentiated for 3 days and then visualised using SiR-Tubulin. Myoblasts cultured on the patterned surface show alignment to the topography, compared to the control conditions where cells are randomly positioned (**Figure 4F**). While the cells follow the given topography, the differences in size and shape of the SOL3D grooves compared to the photolithography pattern, might evoke different interaction and biological responses, making the two different groups not entirely comparable, but Sol3D manufactured grooves provide a suitable alternative to microfabricated grooves and provide guidance and cell alignment in some circumstances. As a resource, the SOL3D grooves provide a suitable alternative to microfabricated grooves and provide guidance and cell alignment. Taken together with the SOL3D stencil-like devices, a whole on-chip platform can be generated using SOL3D.

### 2.4 Customisable SOL3D fabrication as a tailored alternative to standardised commercially available culture platform

The ability to create customisable devices and substrates suitable for cell culture or other biological experiments, with μm to cm sized features, in a fast, reliable and cost-efficient manner would be particularly useful in any wet lab, granting independence from high costs, delivery times and availability of the equivalent commercial products, while enabling substantial customisation. For example, most cell culture vessel layouts are standardised and not tailored to the need of an individual laboratory or a specific cell type, causing higher costs and potential compromises in experimental setups. We therefore aimed to test whether our optimised SOL3D mould protocol could be used to reproduce and further customise relevant features from popular commercially available cell culture products. These constructs can be customised in dimensions and/or shapes for individual experimental aims, while remaining cost-effective, highlighting the versatility and accessibility of our system to enhance biological investigations.

#### PDMS bonding for chamber slide devices

Chamber slide systems and other microscopy-ready hybrid culture devices are commercially available systems that allow cells to be cultured within neighbouring wells directly on cover slides for high resolution imaging, providing small well sizes and imaging-compatible set-ups for high throughput and convenience. To ascertain if our SOL3D protocol could be applied to mimic these constructs, we designed a chamber slide system which can be permanently bonded to an imaging coverslip either using oxygen plasma treatment or a UV sensitive resin adhesive. Importantly, the adaptation of our construct for use with UV resin makes this method accessible to labs without a plasma cleaning system **(Figure 5A).** Our design was fabricated using the SOL3D protocol and was size matched to a 60 mm x 24 mm microscopy coverslip with 12 circular wells with funnel shapes. As described above, we generated a fluidic seal by clamping the device with a glass slide during PDMS curing to isolate neighbouring wells. This extremely flat PDMS surface allows fusion of PDMS to the glass slide using oxygen plasma bonding, or the simple application of a UV adhesive. It is important to note that the UV adhesive is resin based and therefore cytotoxic and cannot be used on any medium-facing area. Astrocyte progenitors were then seeded into selected wells at different concentrations **(Figure 5B).** Staining with a live dye (SiR-Tubulin) revealed an intact fluidic seal in both devices, liquids maintained in the respective wells, and healthy astrocyte progenitor populations. We further demonstrated the high-resolution imaging compatibility (**Figure 5C**), as cells are seeded on a glass slide. Ostensibly, we have demonstrated that both oxygen plasma and UV resin are suitable for PDMS bonding of a chamber slide device and highlighted the capabilities of 3D printing for the fabrication of bespoke chamber slides in a fast and cost-effective way.

**Figure 5:**
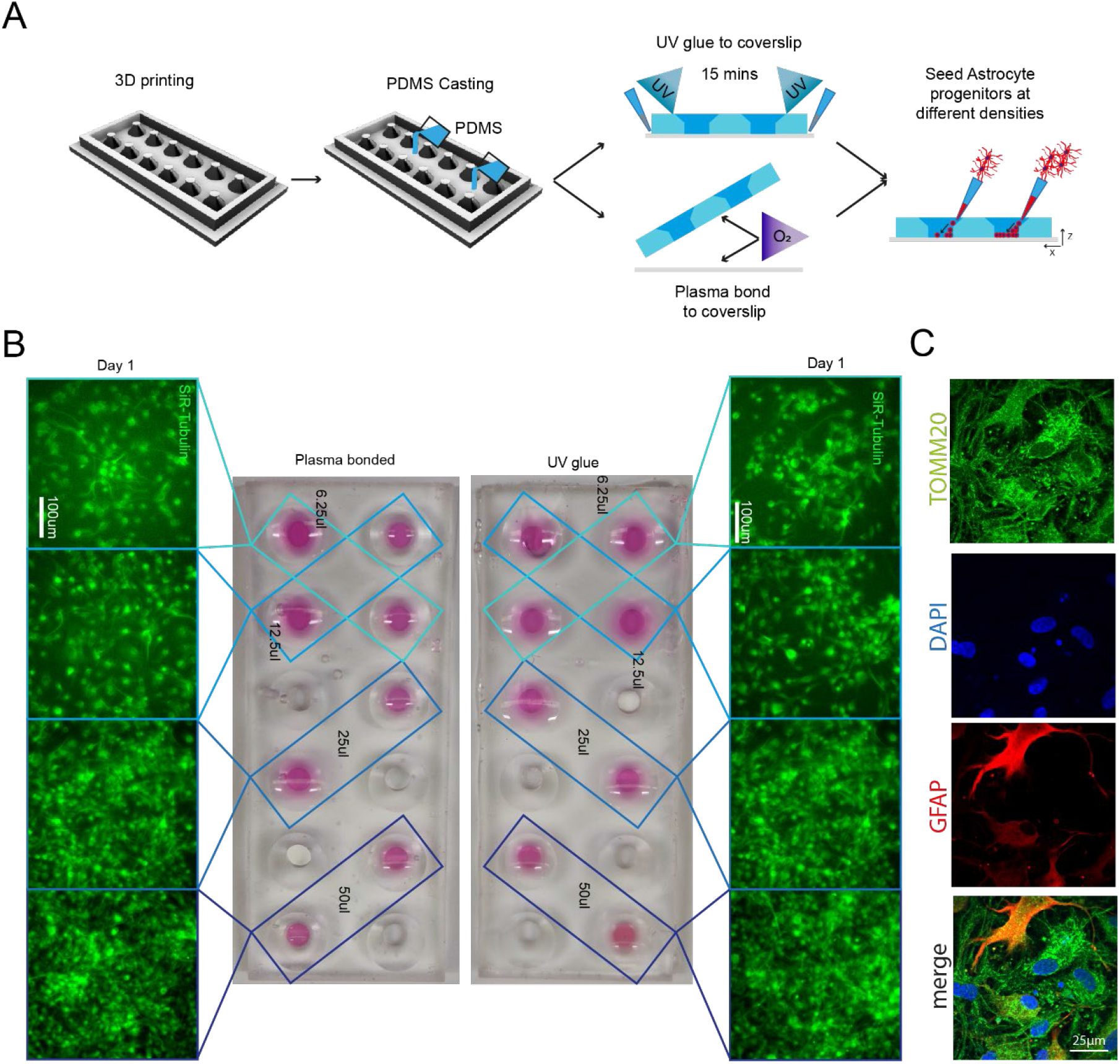
3D printing can create fully customisable imaging chambers with complex well geometry suitable for cell culture via two different methods of PDMS bonding. (A) Schematic overview of design and manufacture of chamber slide device with large wells with (1) UV glue or (2) oxygen plasma bonding to a glass coverslip to seal the wells. (B) As demonstration of the seal quality and viability for cell culture astrocyte progenitors were seeded in different densities in non-adjacent wells and cultured in chamber slide device. Representative images of SiR-tubulin live dye-stained astrocyte progenitors 1 day after seeding in chamber slides bonded with different methods. (C) High-resolution imaging of astrocytes (GFAP) and mitochondria (TOMM20) cultured in custom made SOL3D PDMS chambers.

#### Custom microwell arrays for embryoid body formation

The first design we tested for this purpose was an array of pyramidal-shaped microwells (390 x 350 x 150 µm) which we fabricated using the optimised protocol with no coating step, as it is required for this small feature size (<500 μm) **(Figure 6A-B)**. These microwells have become essential for induction of specific cell lineages from iPSCs and for aggregate research^49, 50^. One of the most important functions of these wells is to ensure homogenous aggregate size for reproducible results, for example, generating embryoid bodies (EBs) of regular size and shape. We used our moulded microwell arrays to form EBs from an iPSCs suspension **(Figure 6C)** and quantified their size after 4 days of culture on the devices. In our microwells, iPSCs formed EBs with consistent diameters, verifying the suitability of our custom PDMS moulds to create small regular arrays of features **(Figure 6D)**. The microwells generated by our protocol are therefore suitable for generation of homogenous EBs with the benefit of substantial customisation of well shape and size at a low cost.

**Figure 6:**
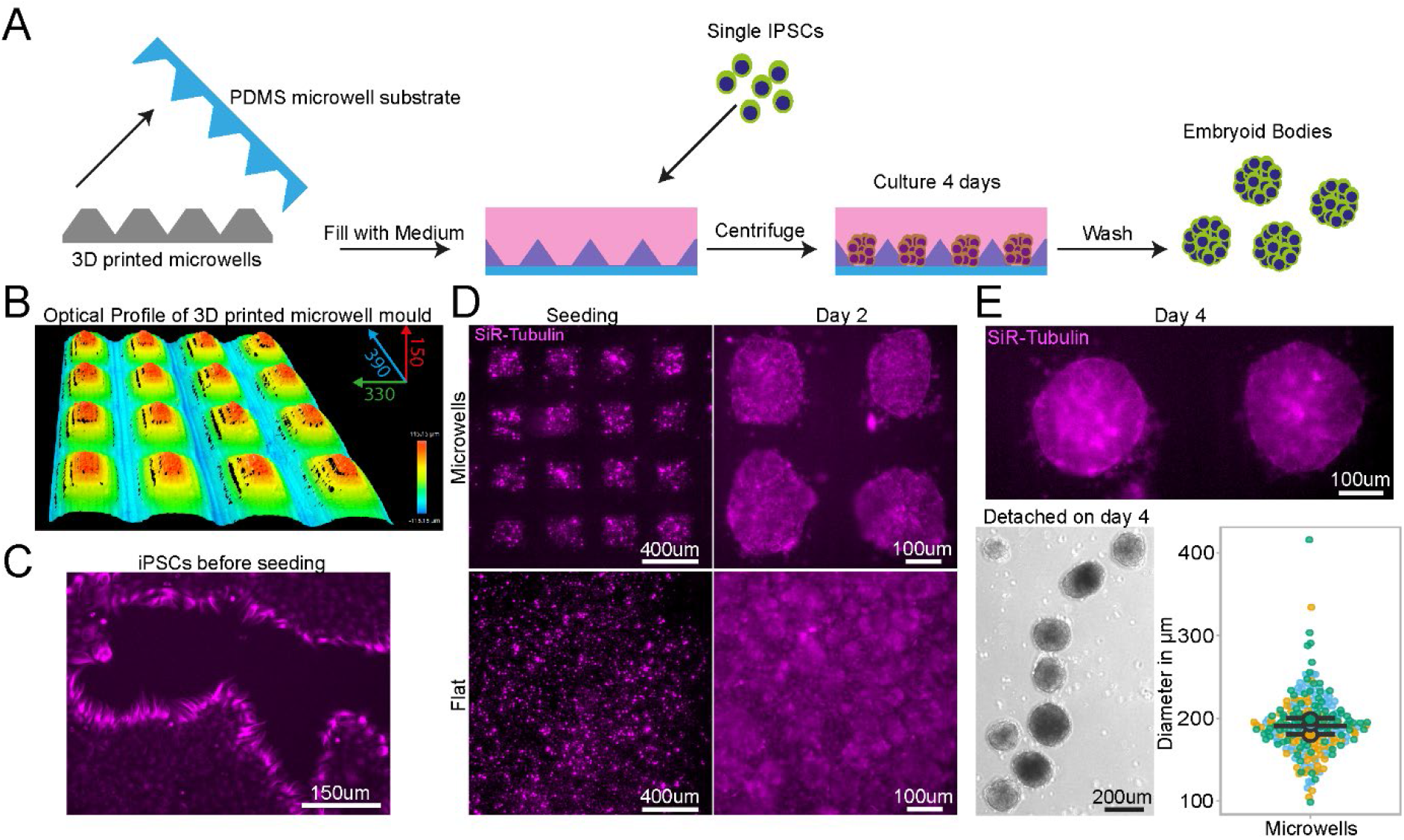
PDMS substrates cast from 3D printed devices permit regular sized embryoid body. (A) Schematic of design, manufacturing and seeding of IPSCs on microwells (B) Representative optical profile of 3D printed microwell device with well sizes of 390 µm length x 350 µm width x 150 µm height. (C) Representative SiR-tubulin live cell dye images of IPSCs before seeding in microwell mould cast. (D) Representative SiR-tubulin live cell dye images of IPSCs seeded on PDMS cast from 3D printed microwell device compared to flat PDMS substrate after seeding (left) and 2 days culture (right). (E) Representative SiR-tubulin image of fused embryoid bodies on microwell PDMS mould prior to detachment after 4 days in culture (top) and embryoid bodies following washing off the PDMS microwell substrate (bottom left). Quantification of embryoid body diameter detached from microwell PDMS mould demonstrates homogenous size of embryoid bodies (bottom right).

#### Large tissue engineered devices with complex designs

Generating complex devices for tissue engineering often combines relatively small features within large constructs and has so far proven challenging to implement in most laboratories, as construction processes are complex and time-consuming, requiring dedicated expertise. Most devices of this kind are therefore sourced from commercially available suppliers, with limited possibility of customisation and at a high cost. For example, tissue-engineered 3D muscle constructs use a variety of devices for suspending large cell laden hydrogels during culture using thin suspension posts ^51, 52^. They are comprised of small pillars with complex shaped end feet, which serve to suspend the hydrogel construct and provide mechanical stiffness to aid differentiation. As these posts are difficult to manufacture and arrive pre-made of a single size and shape, no customisation is available, e.g. miniaturization or altered substrate stiffness. Successful 3D adaptations have been implemented for smaller and less complex muscle post, however with low ease of production for 3D SLA printing ^53^.

We used our SOL3D protocol to fabricate a device for suspended 3D muscle culture with customisable post size and overall dimensions. The challenge, in this case, stems from the fact that these devices do not have large flat faces, they present thin complex features and need ideally to be produced as a single component to avoid complex assembly steps that can introduce variability. A single mould system would in this case not be sufficient, as the lack of air in contact with the complex shapes would prohibit the successful demoulding of the structure. We created a two-part mould/injection system using SOL3D, which can easily be assembled by clamping for curing after PDMS is poured into the mould. Optimisation of the moulds showed that an unequal distribution of the design between the two parts (70/30) is beneficial for successful demoulding (**Figure S15)**, resulting in a reproducible single device with the desired dimensions, in this case twice as large as the commercial alternative – a 2cm muscle compatible with 12-well plates **(Figure 7 A)**. We compared our 12-well plate 3D posts to the commercially available 24-well adapted equivalent (see M&M, *Muscle culture*), using immortalised myoblasts.

**Figure 7:**
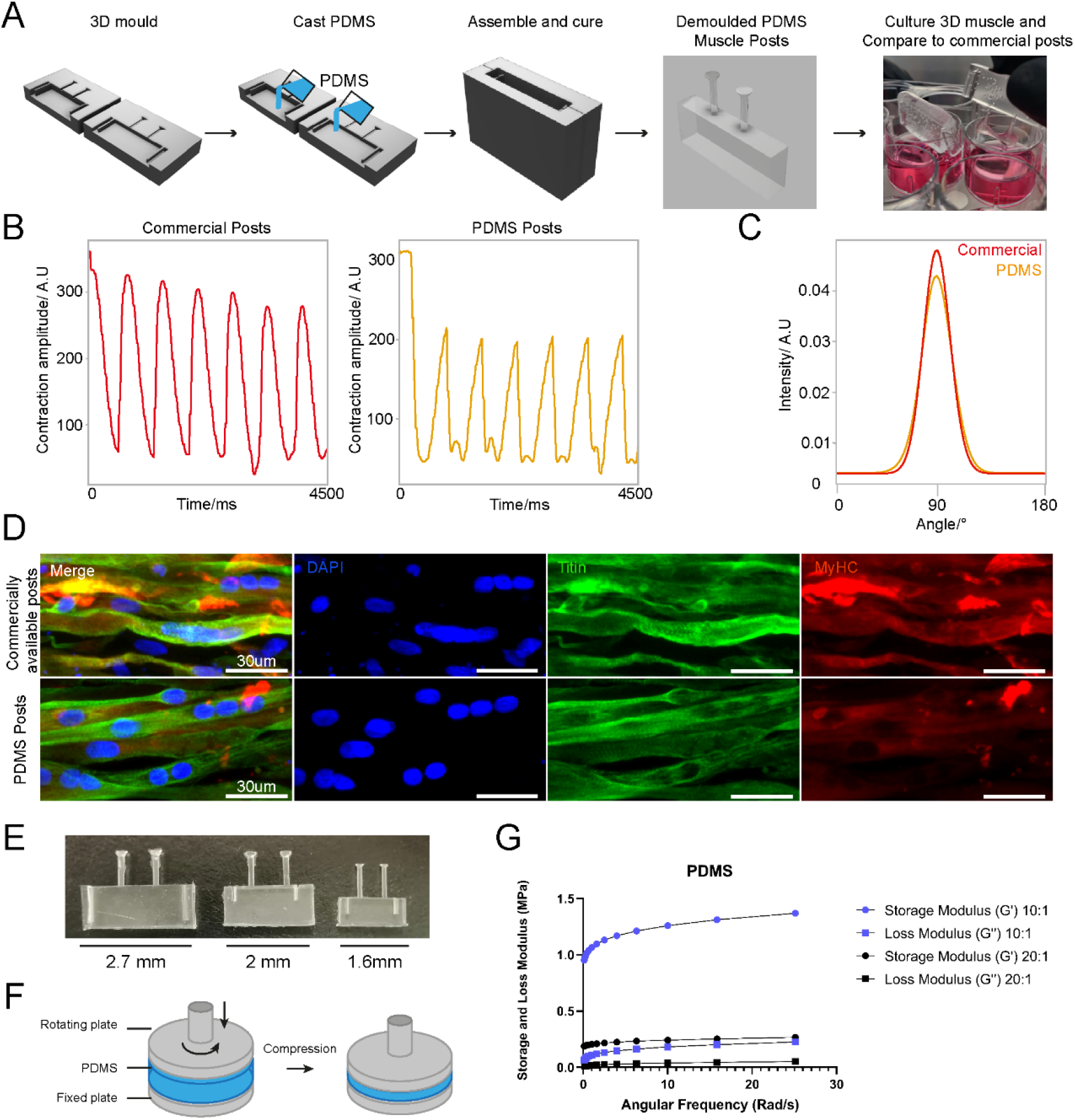
3D sandwich moulds for PDMS casting to generate complex cell culture devices. (A) Schematic overview of design and PDMS casting strategy for a 3D sandwich mould. Immortalised myoblast hydrogels were then formed around PDMS posts, differentiated and cultured for 2 weeks. (B) Comparative contractility analysis with microstimulation at 20mV with 0.5Hz frequency of differentiated 3D muscle between crafted PDMS posts and commercially available posts. (C) Directionality analysis of fiber alignment in differentiated 3D muscle fibers between PDMS posts and commercially available posts after 2 weeks differentiation. (D) Representative images of myoblast differentiation (Titin) and developmental stage (MyHC) on PDMS posts and commercially available posts after 2 weeks differentiation. (E) Images of posts with different dimensions suited for 12, 24 or 48 well plate (from left to right). (F) Illustration of compressing PDMS to measure the modulus through a lateral and rotating movement. (G) The Storage and Loss Modulus of PDMS with different formulations (1:10 or 1:20) at changing angular frequency.

Following the protocol to generate 3D bioengineered muscle constructs described by Maffioletti and co-workers^52^, we first created a pouring mould by filling liquid agarose around a 3D printed rectangular spacer which was removed after the agarose has set (**Figure S9)**. Subsequently, myoblasts were seeded in fibrin hydrogels within the agarose mould, and the SOL3D fabricated posts (or commercially available devices ^54^ used in Maffioletti et al.) were inserted within the still-settling fibrin constructs^52^. After 2 weeks of differentiation, we performed electrical micro stimulation to measure muscle contractility – a hallmark of successful 3D muscle culture-on both constructs at 20mV with 0.5Hz frequency (**Figure 7 B**), which showed periodic contractions for both SOL3D and control devices. Immunostaining of the muscle tissue showed the presence of terminally differentiated myosin heavy chain (MyHC) and titin positive multi-nucleated fibres in both constructs. Directionality analysis revealed that myofibres were preferentially aligned along the posts **(Figure 7 C, D).** After showing the suitability of our 3D posts for muscle cell culture, we wanted to further demonstrate the customisation of such devices. We first focussed on miniaturisation of the posts, which would allow fewer cells to be used and an increased throughput. We designed and manufactured with the SOL3D protocol insets for 12 (i.e. original size) 24 and 48 well plates (Figure 7E). Moreover, as PDMS stiffness can be tuned by changing the ratio between monomer and curing agent, we were also able to produce 24-well plate format muscle devices with different stiffnesses by varying the components ratio (Figure 7 F). As expected, the storage and loss modulus are higher for 1:10 PDMS compared to the 1:20 formulation (Figure 7 G).

In summary, our protocol allows for complex and small features to be easily moulded in PDMS comparable to commercially available 3D muscle systems with the additional benefit of customisation in all aspects of design for improved function.

### 2.5 Entirely SOL3D fabricated hydrogel moulding, culture and imaging system

Hydrogel cultures offer a great opportunity for in vitro 3D modelling, as they are easy to produce, can be tailored to specific applications and provide an *in vivo*-like environment with a complex architecture, and mechanical properties that more align with specific tissues. The low stiffness is a particular advantage for neuronal cultures as the brain is one of the softest matters in the human body, but it also creates challenges in the handling of these hydrogel cultures. Bioprinters can be used to directly form hydrogel cultures with specific shapes and positions of cells within the gel, however, due to the complexity and engineering effort required, accessibility is limited for many labs. Moreover, the use of 3D cultures also presents challenges in adapting conventional culture vessels and plates to the specifics of the hydrogel constructs.

We sought to provide a 3D culture system for hydrogels that allows ease of handling and culture of hydrogels, is entirely customisable and cost-effective, requiring no more than the equipment needed for SOL3D to be implemented.

We started by designing and manufacturing a PDMS mould for shaping hydrogels. This mould is manufactured using the SOL3D protocol and our clamping system for a flat bottom surface so that the mould stays at the desired position. Once placed, pipetting the hydrogel into the mould and seeding of MNs progenitors is performed manually (**Figure 8A**). To further avoid detachment of the gel from the mould and control medium flow to the construct, we developed a chamber system that can be placed on top of the hydrogel with the mould. The design includes a chamber for the hydrogel and mould that is connected with a funnel to a reservoir on the top. This restricts the movements of the hydrogel and at the same time offers the opportunity to provide medium from the top (**Figure 8B**). The chamber has two round-shaped openings that provide a flow of medium from the respective side of the well (**Figure 8C**). We also matched the size of the chamber to the size of a 6-well plate, this allows for providing different media from both sides and the top of the device (**Figure 8 D**). We used IPSC-derived MNs in a custom-made hydrogel and cultured the construct for 7 days (**Figure 8E**). Visualisation of axonal outgrowth in the hydrogel requires fluorescent imaging with a higher magnification. The size of hydrogel constructs limits the simplicity of imaging approaches and usually requires more complicated and expensive setups. We designed an imaging holder which is custom fitted to the size of the hydrogel mould. After staining for axons (β-III-Tubulin), the hydrogel chamber is removed and the mould containing the gel can be easily transferred to the holder. Due to the tailored dimensions, only a small amount of mounting medium is required to cover the sample and then a cover slip is placed on top of the holder (**Figure 8F**). This results in a very tight mounting of the sample, where MNs are close to the glass slide, and minimal disturbance of the sample itself, maintaining the complex architecture (**Figure 8G**). Fluorescent imaging with a low magnification (4x) shows the overall structure of the MN aggregate, and higher resolution images (20x) show that MNs extend long axons in the hydrogel construct (**Figure 8H**). This SOL3D fabricated hydrogel moulding, culturing and imaging system provides an easy and inexpensive system, that is not limited to hydrogels but can be adapted for any 3D system.

**Figure 8:**
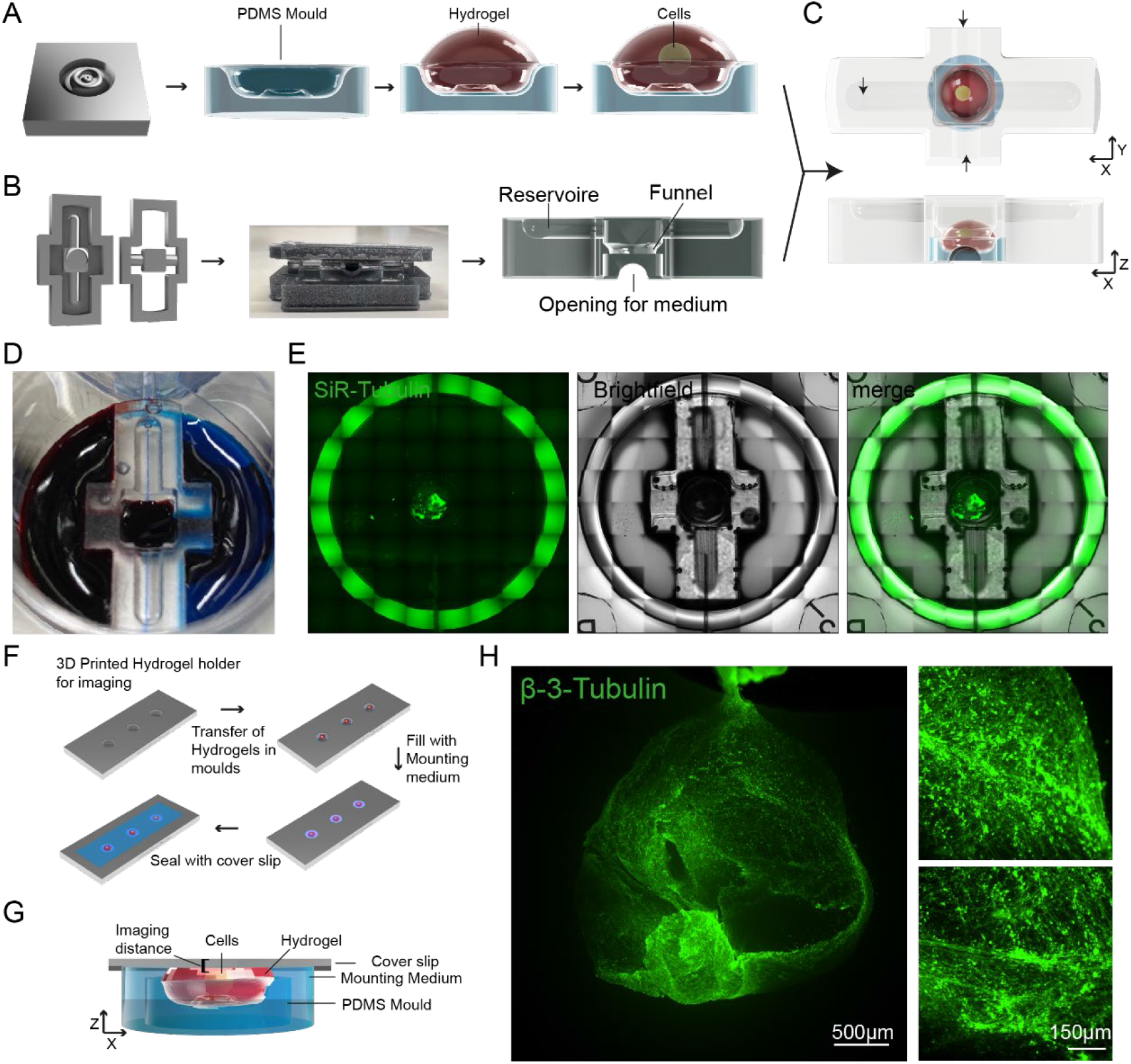
Entirely SOL3D-manufactured hydrogel moulding and culturing chamber system with customised imaging chamber. (A) Schematic overview of PDMS mould generation for hydrogel moulding and MN seeding. Cells are manually pipetted into a preformed hydrogel moulded in the SOL3D fabricated PDMS mould. (B) Schematic overview of the PDMS chamber manufacturing process using SOL3D. The chamber has a complex design with a funnel shape within the structure and multiple openings. We used a two-part design for ease of demoulding. (C) Schematic overview of the combined hydrogel with MNs placed in the mould (A) and a diffusion chamber around the construct (B), arrows indicate the flow of medium. (D) Optional media compartments highlighted using food colouring (blue, red) and PBS (top) (E) Representative images of the SOL3D chamber system (Brightfield) with MNs (SiR-Tubulin) in culture in a 6-well plate. (F) Schematic overview of the mounting process of the complex hydrogel samples for fluorescent imaging. A SOL3D manufactured holder allows the transfer of whole constructs without disturbance and mounting close to the coverslip for imaging. (G) Schematic overview of the hydrogel within the holder. The PDMS mould sits in the bottom of the well with the hydrogel and cells on top. The sample is surrounded by mounting medium and covered with a thin glass coverslip for imaging. Due to the custom holder system, the distance between sample and glass is minimised allowing for fluorescent imaging. (H) Representative fluorescent images of MNs cultured for 7 days in the SOL3D hydrogel chamber system and mounted in the SOL3D chamber. MNs (β-III-Tubulin) seeded in the hydrogel extend many axonal processes.

## 3. Discussion

In this study, we have developed a fast, high resolution and cost-effective protocol to quickly prototype highly versatile cell culture compatible devices that can be applied to a range of different applications, from live cell imaging to microfluidics and even advanced tissue engineering, which we have called **Soft-lithography with 3D Resin moulds using vat polymerisation (SOL3D).** This methodology allows any lab, even those with very minimal prior expertise in the field or without dedicated resources, to effectively set up a complete microfabrication prototyping system and produce culture devices tailored to their specific biological experiments with minimum expenditure.

The widespread commercialisation of 3D printing has led to a significant development of resolution and accessibility, which accompanied by free repositories and software packages, have significantly lowered the entry barrier for the adoption of this technique. As a result of this rapid development and the sheer number of new resins and printers becoming available, it is difficult to get an overview of the suitable protocols and materials for a given application. A general protocol for printing and fabrication using any commercially available product is therefore highly desirable.

We have capitalised on these technological and community developments to overcome one of the primary barriers to complex microfabrication in a biology-focused lab by developing and testing a robust pipeline of fabrication for 3D vat polymerisation, post-processing and PDMS casting, which enables for complete customisation of any cell culture device without further establishment or optimisation. We tested resins from various manufacturers and identified one suitable for PDMS casting and high-resolution prints. In particular, one of the resin compositions performed optimally without the application of a paint layer for high-resolution prints. This feature allows users to exploit the full potential of high-resolution prints, making ink coating for small details redundant^38^. Interestingly, this resin was originally developed for high-resolution printing and has a noticeably lower viscosity than other resins we tested, parameters to consider when evaluating different resin compositions for this method. However, enclosed prints such as the large tissue culture constructs (**Figure 7**) would still not be able to cure sufficiently without a coating layer which isolates the PDMS from the mould. We also tested Resin A with a different commercially available mid-price SL printer and found no observable difference in print quality, PDMS curing, or biocompatibility, showing easy transfer of optimised parameters for resins to multiple printer systems and designs (data not shown).

When considering microfabricated devices for biological experiments, especially with cell culture and other *in vitro* set-ups, the topographical features one might want to add can range roughly from sub-cellular scale (<5 μm, e.g. nanoindentations^55^ and other nanostructures^56^), cellular scale (10-100μm e.g. microgrooves^57^ and other microwells^58^), multicellular (200-1000μm e.g. microfluidic channels^59^) or tissue scale (>1mm e.g. aggregate culture devices^60^). In the vast majority of microfabrication pipelines, there is a practical gap at the interface between the cellular and multicellular scale, as conventional photolithography is mostly suited for precise features on the smaller scales and is less suited for multicellular scales. Moreover, combining efficiently multiple fabrication rounds across scales is sometimes challenging, time consuming and error prone. Instead, methodologies such as SOL3D are capable of simultaneously combining cellular, multicellular and tissue scale features within the same fabrication round, effectively filling a gap in the potential toolkits currently available.

Another advantage here is that SOL3D is not tied to any particular printer or resin, and as such can work for any combination (we have tested several resins over 3 different printers) and with the advent of higher resolution printers this gap will become significantly smaller. We have also demonstrated the versatility of our pipeline by successfully developing cell culture devices of different designs targeted to a wide range of applications, which either confers new capabilities to conventional culture systems (e.g. easily plating multiple cell types on precise spatiotemporal relationships) or customisation of bioengineered culture systems.

While a number of studies have already proposed similar protocols (e.g. heat curing^61^, UV light^62^, micro-diamond coating^63^), they all generally tend to either have a considerably lower resolution or in some cases require obligatory steps with very expensive specialised equipment that is not generally available to most biological labs. Moreover, the chemicals and materials used in several of these processes require a much higher degree of training and expertise compared to the pipeline presented here, which requires no hazardous processes or chemicals, but only non-toxic and easily handleable components, and could therefore be implemented in a lab without risk.

In its most minimal implementation, SOL3D requires only a high-resolution (∼50µm) desktop vat polymerisation printer, a suitable resin, PDMS and everyday cell culture and microscopy components. All of these resources can be obtained with an initial investment below the cost of 2 vials of monoclonal antibodies and at an estimated running cost of <300USD per year to produce regular medium size batches of devices, which would be affordable to any lab that has an ongoing budget for cell culture consumables. For context, a commercially available single-use 8-well chamber slide suitable for live microscopy with fixed dimensions and no possibility of customisation, ranges between 10 and 15 USD, while a single microfluidic device costs from 50 to 120 USD, both of which can be easily replaced by SOL3D.

One caveat to the capability of the method, which is a limitation due to the properties of PDMS rather than the fabrication process, is that while it is possible to create complex devices without any specialised equipment to effectively use smaller features (**Figure 2**), a step of oxygen plasma treatment was necessary. However, plasma etchers are not available to all labs, potentially limiting the applicability of this protocol. Wang and colleagues tested alternatives to plasma treatment of PDMS persorption of fibronectin sufficient for Caco-2 cells^64^, hence plasma treatment might not be necessary for all cell types and has to be assessed individually.

Unfortunately, the low compatibility of most UV resins with cell culture experiments also poses limitations to the potential application of this method and vat polymerisation in general. We have characterised several different commercially available resins and found that all of them, in a treated or untreated state, are cytotoxic (**Figure S1**). There are some biocompatible alternatives available on the market (although the extent to which biocompatibility, as defined for dental implants, can be directly applied to stem cell cultures and other more sensitive biological systems needs to be verified) to overcome this fundamental limitation of 3D vat polymerisation, but for a far higher price than any standard resin. Additionally, the fact that resin composition is generally proprietary limits the possible customisations that end-users can achieve. Some providers have recently started publishing their resin composition as well as customised resins generated from a number of research groups that can be recreated, although the burden of adoption of these unique formulations is likely to prevent their use in most cases^65–67^. Alternatively, modifications of PDMS to be printable can lead to decreased resolution of the print, undermining the strong advantage of high-resolution printers for microfabrication using resins^68^. The solution we implemented for the well-known curing inhibition issues with resin moulds and PDMS is to extract and cure completely the resin first with a multistep preparation (see **Figure 1**) and then further shield the PDMS with a layer of enamel coating.

The use of PDMS in cell culture experiments has a long history and we have decades of experience with its use with cells and in microscopy applications, however it also has some potential drawbacks. In particular, several groups have reported that PDMS can retain organic molecules and adsorb other substances, which can severely impact biological and biochemical experiments in particular cases ^35, 69–71^. However, one advantage of this setup is that it can be used for any soft-lithography material, for example, polymers such as Flexdym, polymethylmetacrylate, or poly(DL-lactide-co-glycolide), as well as with any hydrogel or cure-forming material with a temperature below 70 degrees.

With this system we aim to empower any lab, regardless of its capabilities, access to resources or prior expertise, to create customised microdevices with features tailored to their specific biological experiments and designed with their biological question in mind, which will significantly lower the barrier for experimentation with microfabrication and tissue engineering applications in any field.

## 4. Materials & Methods

### Cell culture

Control hiPSC motor neuron and cortical neuron progenitors were derived as described^72^ from multiple donors (Table 2). These cells were cultured on Matrigel (Corning) coated plates in base medium, comprised of 50% NeuroBasal (Gibco), 50% advanced DMEM (Gibco), supplemented with B27 and N2 (gibco),100 µg/ml Pen-Strep (Gibco), and 2mM L-alanyl-L-glutamine dipeptide (Gibco). For expansion of progenitors, FGF (20 ng/ml) (Gibco) was added to base medium. Differentiation of MN progenitors was achieved using base medium with compound E (Enzo) (0.1 uM) and the growth factors BDNF (10 ng/ml) and GDNF (10 ng/ml) unless stated. Astrocytes were generated from iPSC using a modified protocol described in Hall et al^72^. Derived astrocyte progenitors were cultured in neuronal base medium with FGF (20 ng/ml) (Gibco) and EGF (20 ng/ml) (Thermo). For differentiation astrocytes were cultured in base medium without growth factor supplements. All cells were cultured with 5% CO2 in humidified atmosphere at 37°C (Table 1).

**Table 1:**
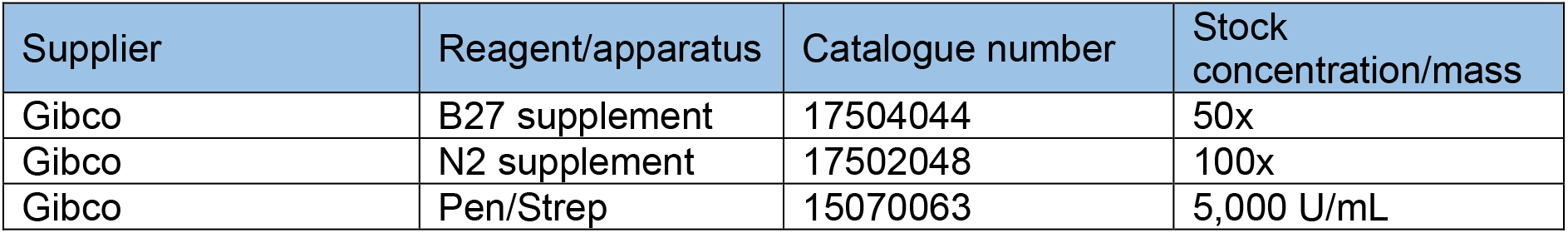
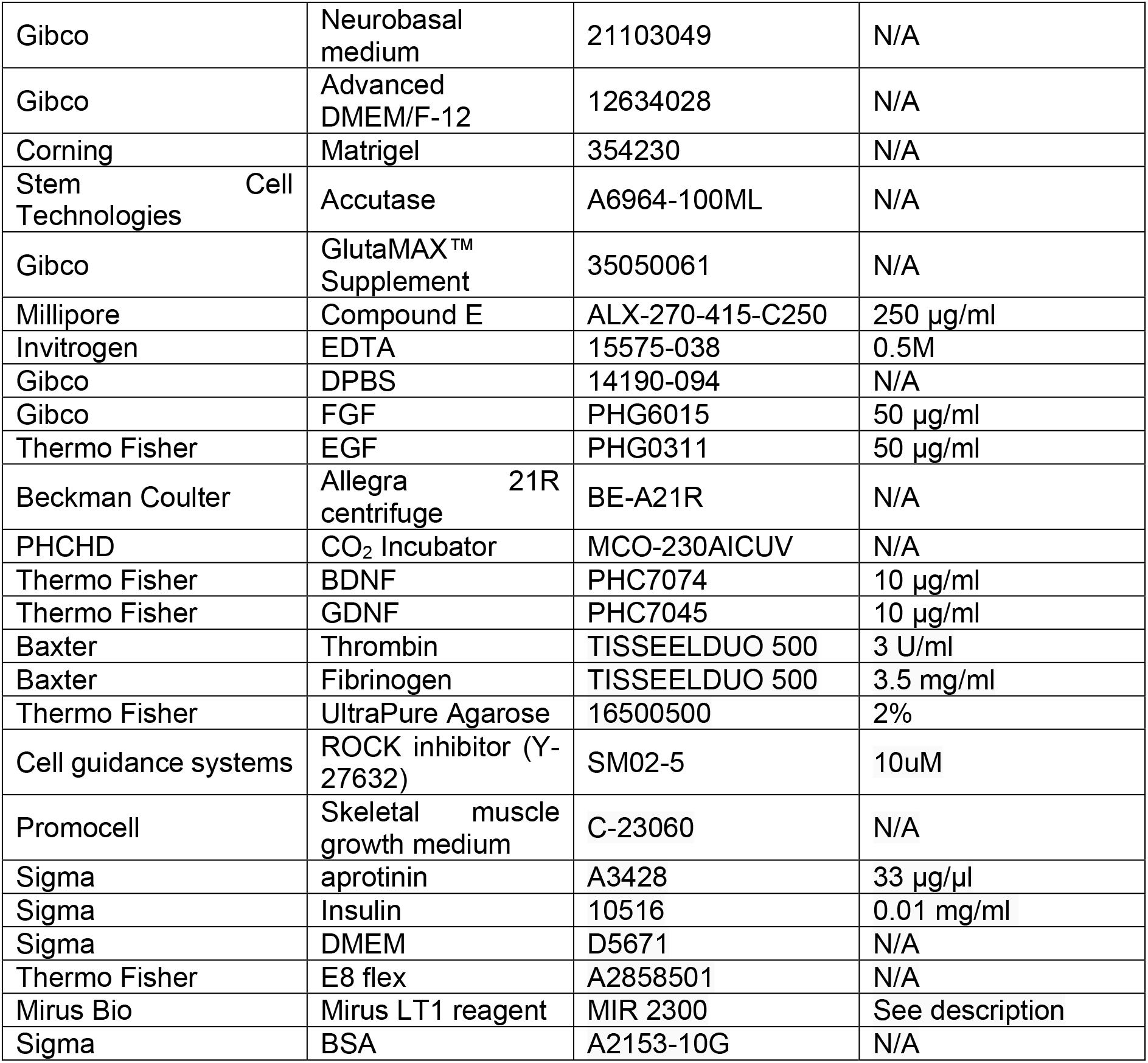
Cell culture

**Table 2:** Cell lines

| Cell lines | Name | Donor Age | Donor Sex | Mutations present |
| --- | --- | --- | --- | --- |
| iPSC/ Motor neuron/ cortical/ astrocyte progenitors | Control 1 | 78 | Male | None |
| iPSC/ Motor neuron/ cortical/ astrocyte progenitors | Control 2 | 64 | Male | None |
| iPSC/ Motor neuron/ cortical/ astrocyte progenitors | Control 3 | unknown | Female | None |
| iPSC/ Motor neuron/<br>cortical/ astrocyte<br>Progenitors | Control 5 | 51 | Male | None |
| AB1190 Human<br>immortalised<br>myoblasts | Paravertebral<br>muscle | 16 | Male | None |

### Transfection of MNPs

For transfection of cells a plasmid based *Piggybac* transposon system was used with pgK-Puro-CMV-GFP and pPb-CAG-RFP-Hygro construct cloned in the lab and a *PiggyBac* vector containing the transposase. Motor neuron progenitors were sparsely cultured on a 24-well plate and one day after passaging transfected using Mirus LT1 (Mirus Bio) transfection reagent. Plasmids were added at total 0.5 µg per well (GFP/RFP+ transposase containing plasmid) in 200 µl of Pen-Strep free growth medium. This solution was gently mixed before addition to the wells which also contained 200 µl of Pen-Strep free medium. The medium containing the transfection reagent was exchanged with growth medium after 24h. Cells were cultured to confluency and then pooled into a 6-well plate for further expansion.

### Fabrication

#### 3D printing

3D printed moulds were designed using fusion 360^73^ and tinkercad^74^ computer aided design software, then exported as .stl (stereolithography) files to either Chitubox or Photon workshop slicing software. These software were used to define print parameters such as layer thickness, layer UV exposure time, and lifting/retract speeds for each resin. All resins, printers and print settings can be found in Table 3 and Table 4.

**Table 3:** Resins

| Supplier | Resin | Synonym |
| --- | --- | --- |
| Phrozen | Aqua Gray 4K | A |
| Elegoo | ABS-like | B |
| Anycubic | Clear | C |
| Liqcreate | Flexible X | D |
| Liqcreate | Premium Tough | E |
| Phrozen | Water-washable model Gray | F |
| NextDent | Ortho Clear |  |

**Table 4:** 3D Printer Fabrication

| Supplier | Resin | Printer | Layer thickness<br>($\mu\text{m}$) | Layer exposure<br>time (s) | Bottom<br>Exposure time<br>(s) | Bottom Layers | Transition<br>layers | Light off delay<br>(s) | Z Lift Distance<br>(mm) | Lift<br>speed<br>(mm/min) | Retract<br>speed<br>(mm/min) |
| --- | --- | --- | --- | --- | --- | --- | --- | --- | --- | --- | --- |
| Phrozen | Aqua<br>Gray 4K | Phrozen<br>sonic<br>mini 4K | 50 | 6.0 | 50 | 6 | 0 | 9 | 5 | 65 | 150 |
| Phrozen | Water-washable<br>model<br>Gray | Phrozen<br>sonic<br>mini 4K | 50 | 2.5 | 25 | 4 | 0 | 9 | 5 | 65 | 150 |
| Elegoo | ABS-like | Anycubic<br>Photon S | 50 | 9.0 | 60 | 6 | 0 | 1.5 | 6 | 180 | 180 |
| Anycubic | Clear | Anycubic<br>Photon S | 50 | 10.0 | 50 | 8 | 0 | 1 | 6 | 180 | 180 |
| Liqcreate | Premium<br>Tough | Phrozen<br>sonic<br>mini 4K | 50 | 6.0 | 75 | 1 | 6 | 9 | 5 | 65 | 150 |
| Liqcreate | Flexible<br>X | Phrozen<br>sonic<br>mini 4K | 50 | 18.0 | 70 | 1 | 6 | 9 | 5 | 65 | 150 |
| Phrozen | Aqua<br>Gray 4K | Phrozen<br>sonic<br>mini 4K | 10 | 1.8 | 37.5 | 6 | 0 | 10 | 5 | 55 | 150 |
| Next Dent | Ortho Clear | Phrozen sonic mini 4K | 50 | 4.6 | 30 | 6 | 6 | 10 | 6 | 60 | 150 |

#### Post processing

After printing, constructs were washed in fresh isopropanol (IPA) using either/or/both an ultrasonic cleaner and stirring washing bath (Anycubic). Washing method and time were varied as part of the protocol establishment. To ensure fair comparison, washing IPA was filtered for every resin for a given washing condition to remove resin components from previous washes. After washing, all prints were cured in a commercially available curing chamber (Anycubic) for 60 mins. Constructs were then selectively coated with a layer of enamel paint (Plastikote) using a hobbyist airbrushing system (Timbertech) diluted 70:30 with water as per manufacture instructions. Painted casts were left to dry at room temperature on the bench for at least an hour before PDMS casting (Table 5).

**Table 5:**
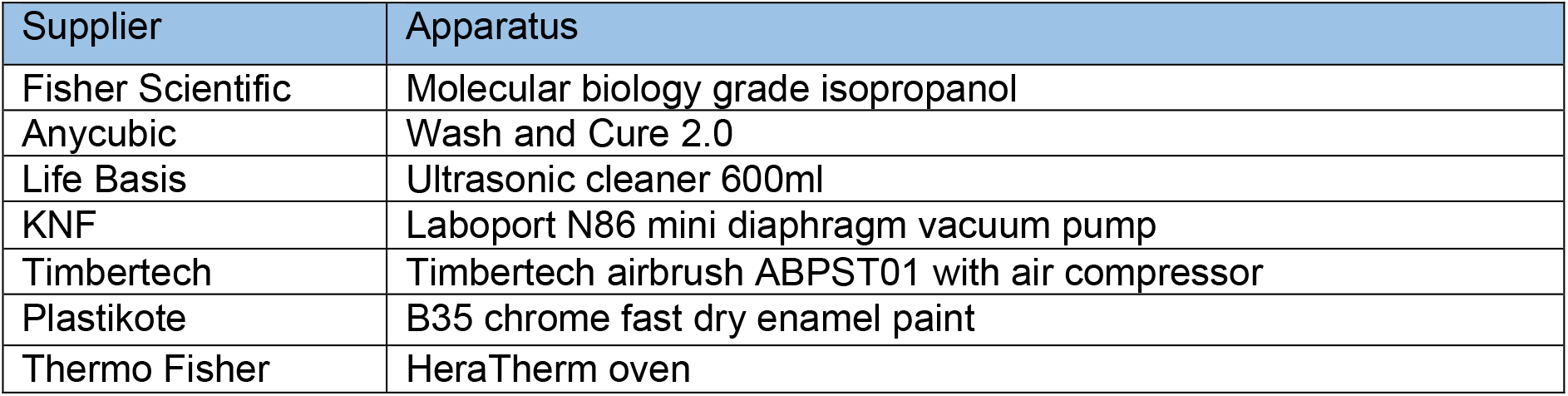
Post Processing

#### Microfabrication of patterned substrates

Microgroove substrates were manufactured from silicon masters patterned using photolithography as previously described ^75^. Briefly, SU-8 2002 (Kayaku) was spun on a silicon wafer for 40s at 1000 rpm on a spin coater (Polos) and prebaked at 95 °C for 2 minutes. A microgroove pattern designed in CleWin5 and containing 10x10 µm grooves with 250 µm plateaus every 5 mm was then etched into the SU-8 via UV exposure and an aligned photomask with the design (Kiss MA6 mask aligner). Excess SU-8 was cleaned with PEGMA, then soft and hard baked at 95°C for 5 mins, before being silanized with trichlorosilane (C_8_H_4_Cl_3_F_13_Si) in a vacuum chamber for 1 hour. Excess silane was then washed off from masters with 100% acetone. An unpatterned silicon wafer was used for flat substrates. For comparison of surface roughness between 3D printed casts and microfabricated substrates, etching was achieved by a single photolithographic step using a MicroWriter ML3 (Durham Magneto Optics) to form a pattern designed to mimic the potential capabilities of 3D printing in microfabrication. Following the photolithographic step, PDMS casts (prepared as above) were made of the flat or micropatterned silicon wafers, spin coated at 300 rpm for 40s on a spin coater (Polos) to ensure uniform thickness and cured on a hotplate at 100 °C for 5-10 minutes (Table 6).

**Table 6:** Microfabrication

| Supplier | Reagent/Apparatus | Catalogue code |
| --- | --- | --- |
| Kayaku | SU8 - 2002 | N/A |
| Karl Suss | MA6 mask aligner | N/A |
| Sigma Aldrich | Trichlorosilane | 448931 |
| Sigma Aldrich | PEGMA | 409537 |
| Fisher Scientific | Acetone | 13277983 |
| Fisher Scientific | Vacuum Desiccator | 11852732 |
| Durham Magneto Optics | MicroWriter ML3 | N/A |
| Polos | 200 NPP Spincoater | 42839 |

#### PDMS

Sylgard-184™ silicone elastomer kit PDMS pre-polymer was well mixed (5 mins) with curing agent at a 10:1 w/w ratio using a digital balance (Sartorius BP610) prior to vacuum desiccation and casting at various temperatures (60-90°C).

#### Biofunctionalisation

Biofunctionalisation of PDMS substrates and casts from 3D printed moulds was achieved using oxygen plasma treatment (30s, 50%, 7sccm – unless stated otherwise) (Henniker Plasma). Additional biofunctionalisation of micropatterned PDMS substrates to facilitate cellular attachment was achieved using a coat of poly-D-Lysine 0.01% (PDL) (Gibco) for 15 min and laminin (Sigma) overnight (unless stated otherwise) (Table 7).

**Table 7:**
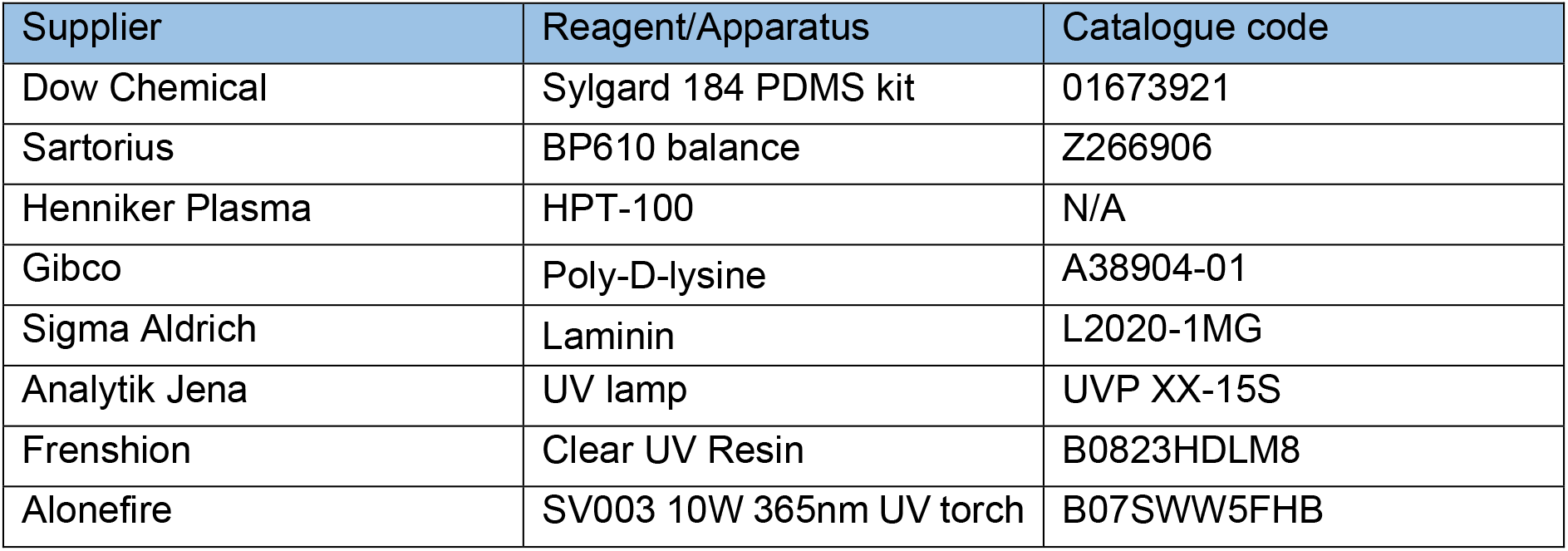
PDMS soft lithography and biofunctionalisation

**Table 8:** Surface characterisation

| Supplier | Reagent/Apparatus | Catalogue code |
| --- | --- | --- |
| Phenom | ProX Desktop SEM | N/A |
| Sensofar | S Neox optical profiler | N/A |
| Quorum | Q150R coater | N/A |

**Table 9:** Immunocytochemistry

| Supplier | Reagent | Catalogue code |
| --- | --- | --- |
| Sigma Aldrich | Goat serum | G9023-10ML |
| Invitrogen | 1% Triton in PBS | HFH10 |
| Millipore | Fluor Save | 345789 – 20 ml |
| Fisher Scientific | 1 mm microscope slides | 15545650 |
| Fisher Scientific | 22x22 glass coverslips | 12312128 |
| Fisher Scientific | 60x24 glass coverslips | 10083957 |
| Boster | 4% Paraformaldehyde | AR1068 (500 ml) |
| Sigma Aldrich | Brand™ Cavity Slides | BR475535-50EA |
| Thermo Fisher | Fluoromount G | 00-4958-02 |

### Cell Seeding

Cells were detached from culture using Accutase (Stem Cell Technologies) and seeded in oxygen plasma treated PDMS constructs cast from 3D printed moulds bound to either tissue culture plastic, flat PDMS, or micropatterned PDMS substrates. Cells were concentrated to 300 µl per detached well and seeded in differentiation media with Compound E (Milipore). Following initial plating, cells were left to settle for 2 hours in constructs. Cells were then washed 2x with PBS to remove unattached cells from constructs and wells filled with differentiation media supplemented with Compound E with constructs left in place. After 24 hours in culture, cells were washed with PBS and constructs removed before a 1:100 Matrigel spike for >2 hours and culturing cells (as above) in differentiation media supplemented with Compound E for 7 days (or as stated). For longer term experiments media was selectively supplemented with additional BDNF and GDNF growth factors depending on the experiment.

### Immunostaining

Prior to staining cells were fixed in 4% PFA (Boster) for 15 min (unless stated otherwise) and washed 3x with PBS. Cells were then permeabilized with 0.1% Triton-X for 10 min and blocked with 3% goat serum (GS) (Sigma) for 30 mins at room temperature (unless stated otherwise). Antibodies diluted in 0.05% Triton-X and 1.5% GS in PBS were then used to stain cells for markers of interest. Antibodies and their concentrations can be found in Table 10. Primary antibodies were incubated for 1 hour at room temperature in the dark, before washing 3x with PBS. Secondary antibodies were then incubated for 30 min at room temperature in the dark, before washing 3x with PBS. All secondary antibodies were incubated at a final concentration of 2 µg/ml (Table 11). For some experiments, stained cells were then mounted on glass slides using FluorSave (Millipore). Otherwise, cells were kept in PBS at 4°C until imaging. For live cell imaging cells were incubated with 1:10,000 (100nM) silicon rhodamine tubulin (SiR) live cell dye (Spirochrome) for 1 hour before removal of the dye and imaging.

**Table 10:** Primary antibodies/Stains

| Supplier | Antigen | Species | Isotype | Clone | Catalogue Code | Dilution |
| --- | --- | --- | --- | --- | --- | --- |
| Sigma Aldrich | βIII-Tubulin | Mouse | IgG2b | SDL.3D10 | T5076 | 1:1000 |
| Thermo Fisher | MAP2 | Mouse | IgG1 | M13 | 13-1500 | 1:500 |
| Abcam | GFAP | Chicken | Polyclonal | Polyclonal | ab4674 | 1:500 |
| Thermo Fisher | DAPI | N/A | N/A | N/A | 62248 | 1:5000 |
| Spirochrome | Silicone Rhodamine-tubulin kit | N/A | N/A | N/A | SC002 | 1:10,000 |
| DSHB | MF-20 | Mouse | IgG2b | MYH1E | MF-20 | 1:9 |
| DSHB | Titin | Mouse | IgM | TTN | 9 D10 | 1:20 |

**Table 11:** Secondary Antibodies

| Supplier | Host species | Target species | Isotype | Conjugate | Catalogue Code | Dilution |
| --- | --- | --- | --- | --- | --- | --- |
| Invitrogen | Goat | Mouse | IgG2b | Alexa Fluor 647 | A-21242 | 1:1000 |
| Invitrogen | Goat | Mouse | IgG1 | Alexa Fluor 555 | A-21127 | 1:1000 |
| Invitrogen | Goat | Chicken | Polyclonal | Alexa Fluor 555 (H+L) | A-21437 | 1:1000 |
| Invitrogen | Goat | Mouse | IgG2b | Alexa Fluor 546 | A-21143 | 1:1000 |
| Invitrogen | Goat | Mouse | Polyclonal | Alexa Fluor 488 | A-21042 | 1:1000 |

**Table 12:** Microscope hardware

| Supplier | Apparatus | Catalogue code |
| --- | --- | --- |
| Nikon | TE-2000-E | N/A |
| Solent Scientific | CAL3300 incubator temperature regulator | N/A |
| Cool LED | pE-4000 16 LED light source | N/A |
| Okolab | CO<sub>2</sub> regulator | N/A |
| Zeiss | LSM710 Inverted Confocal microscope | N/A |

**Table 13:** Microstimulation Hardware

| Supplier | Reagent/Apparatus | Catalogue code |
| --- | --- | --- |
| EHT Technologies | Silicone posts | C0001 |
| EHT Technologies | Carbon pacing electrodes | P0001 |
| DinoLite | USB microscope | AM73915MZT |

### Microscopy

Cells were imaged using an encased Nikon eclipse TE2000-E fluorescence microscope running Micromanager software with 4x, 20x LWD and 20x SWD objectives, cool LED pE-4000 16 LED light source, and a Prior controlled stage. An LED driver Arduino controlled light source provided illumination for brightfield imaging. To allow longitudinal and live imaging, the microscope chamber was humidified and heated to 37.0°C with 5% CO_2_ using a CAL3300 incubator temperature regulator (Solent Scientific) and CO_2_ regulator (Okolab). Humidity, CO_2_ balance, and temperature were regulated by further encasing the plate in a sealed custom 3D printed chamber with humidified CO_2_ inlet.

### Surface Characterisation

#### Optical profilometry

Quantification of 3D printed mould dimensions and surface roughness was achieved using a Sensofar S Neox optical profilometer to measure features in X and Y, and layer thickness. Multi-image z-stacks were captured over stitching areas with 25% overlap using a 20x Nikon EPI objective and surface-variation scanning mode. For analysis of patterned silicon master feature dimensions and PDMS cast surface roughness from both 3D printed moulds and silicon masters, multi-image z-stacks were captured of stitching areas with 25% overlap using a 20x Nikon DI objective and confocal scanning mode. Analysis of features was conducted using in-built analysis tools. Plane correction was conducted on all images to reduce bias within imaged ROIs.

#### SEM

Samples of coated and uncoated 3D printed moulds were sputter coated with a 10 nm thick layer of Platinum using a Quorum Q150R coater and imaged by a Phenom ProX Desktop SEM (Thermo Scientific) at an acceleration voltage of 10kV (unless noise was too high, then 5kV was used). Images of the surface and cross section of prints was captured to investigate the thickness of applied paint and identify changes in surface roughness/topography. Images were processed in ImageJ FIJI ^76^(Table 8).

### Image analysis

All image analysis was conducted in ImageJ FIJI software, processed using R, and presented with R or super-plots-of-data app ^77^ unless stated otherwise.

#### Line Graph Analysis

To quantify cellular segregation within the same device/multiple devices threshold fluorescence intensity was adapted to improve signal-to-noise ratio. A rectangular area was then drawn over the cells of interest and fluorescence intensity plots were obtained for each point.

#### Area and aspect ratio

Measurements of seeded cell area and aspect ratios were compared to either CAD specifications or 3D printed mould feature dimensions. Cell measurements were taken from the borders of aggregates using tubulin markers Silicon-Rhodamine tubulin (live) and βIII-tubulin (fixed). 3D printed mould dimensions were obtained using SensoScan software in-built analysis tools.

### Resin biocompatibility

Chips from each of the 6 resins printed using 50 µm layer thickness printer settings (Table 4) and post processed as above (20 mins sonication & wash, 60 mins UV cure) without enamel coating. Chips were then either sterilised with UV for 15 mins, or baked for 4 hours at 75°C, washed in PBS for 72 hours at 50°C, and UV sterilized for 15 mins before being added to cultures of motor neuron progenitors pre-stained with SiR-tubulin cytoskeletal live dye. Cells incubated with untreated chips were left for 48 hours before imaging. Cells incubated with treated resin chips were live imaged for the first 24 hours in culture, then again at 48 hours with control wells not containing any resin. Investigation of the Ortho dental clear resin, was performed on cylinder shaped resin prints, that can be used as an inset to tissue culture wells.

The resin prints were either treated using our SOL3D protocol or the washing protocol suggested by the manufacturer. The processed resin tubes were then inserted into wells containing a known number of iPSC-derived MNs and coculturerd for 4 days.

### Microwell arrays

Microwell arrays of repeating 400 µm x 400 µm x 150 µm pyramids were 3D printed at 10 µm layer thickness (Table 4) and post processed as above (20 mins sonication & wash, 60 mins UV cure) without enamel coating. PDMS (prepared as above) was casted and left un-biofunctionalised to enhance EB formation through PDMS hydrophobicity. Single IPSCs pre-stained with SiR-Tubulin live dye were then seeded on microwell arrays or flat PDMS in E8 flex medium (Thermo) and centrifuged for 1 min at 100 rpm to settle cells into microwells. Cells were cultured for 4 days, imaged directly after seeding, and at day 2 and 4 of culture. After imaging on day 4 the regular EBs were detached from microwells and imaged to measure their size.

### Chamber slide

#### Manufacture

A 3D design with 12 wells with 5 mm diameter funnel and 3 mm diameter 1 mm deep straight well was 3D printed with Resin A at 50 µm layer thickness (Table 3) on the Phrozen Sonic mini 4K and post processed as above (20 mins sonication & wash, 60 mins UV cure) with enamel coating. PDMS (prepared as above) casts were made of the arrays and bonded to glass coverslips with two methods.

#### PDMS bonding Oxygen Plasma

The surface of PDMS casts from 3D printed moulds and glass coverslips were oxygen plasma treated (30s, 50%, 7sccm) (Henniker Plasma), sealed together, and baked at 75°C for 15 mins.

#### UV glue

The surface of PDMS casts from 3D printed moulds was sealed on glass coverslips and clear photopolymerisable resin was applied round the exterior of the PDMS. Through visual inspection we verified that the glue is only applied on the outside of the device and cannot physically leak into the wells. This was verified though live cell imaging on the chamber device, a uv resin is highly toxic for cells and would have resulted in immediate cell death. Resin was cured via 1 minute UV exposure with a 365nm UV torch (Alonefire).

#### Biofunctionalisation

Plasma and UV bonded PDMS chamber slide devices were then sterilised with UV for 15 mins (Analytik Jena UV light) and biofunctionalized with a coat of poly-D-Lysine 0.01% (PDL) (Gibco) for 15 min and laminin (Sigma) overnight in each well.

#### Seeding

Control 3 astrocyte progenitors pre-stained with SiR-tubulin live dye were then seeded (as defined in methods section – Cell Seeding) after concentration (1 confluent well in 400 μl media) in expansion media with decreasing density in alternate wells of the chamber slide device. Cells were seeded in 50 µl, 25 µl, 12.5 µl, 6.25 µl fractions of the concentrated cell stock, left to settle for 10 mins, before additional media was added. Cells were cultured as before (Methods – Cell Culture) and imaged after 24 hours to qualitatively assess viability at different densities.

### 3D muscle

#### Preparation, differentiation and culture

3D muscle constructs were prepared as previously described^52, 78^. Rock inhibitor (Cell guidance systems) at a concentration of 10µM was added to cells 2 hours prior to preparing gels. Per construct 1 x 106 AB1190 myoblast cells were used in a total volume of 120 µl comprised of 3.5 mg/ml of human fibrinogen (Baxter, TISSEELDUO 500), 3 U/ml of thrombin (Baxter, TISSEELDUO 500), 10% Matrigel (Corning, 356231), and inactivated myoblast medium (20-30 mins at 56°C) (PromoCell skeletal muscle cell growth medium, C-23060). The mix was pipetted into agarose moulds containing posts and placed at 37°C, 5% CO_2_ for 2 hours allowing hydrogels to polymerise. These moulds were prepared in 12-well plates using 2% UltraPure agarose (ThermoFisher Scientific, 16500500) (**Figure S8B**). A ring was placed underneath the arms of the posts and inserted onto a 12-well plate to determine the distance the posts can be pushed down into the agarose (**Figure S8C**). DMEM (Sigma, D5671) was then added to the construct and the posts containing the muscle construct was removed from the agarose mould and placed into a new 12-well plate with myoblast medium containing 33 µg/µl aprotinin (Sigma, A3428) at 37°C, 5% CO_2_. After 48 hours, the muscle construct was placed in differentiation media (DMEM with 0.01 mg/ml insulin (Sigma, 10516) and 33 µg/µl aprotinin), media was changed every other day.

#### Contractility analysis

Artificial muscle contractility on 3D print moulded PDMS and commercially available posts^54^ was achieved via microstimulation using an in-house stimulator system with a pair of autoclaved pacing carbon electrodes (EHT-technology) mounted in the well containing the 3D muscles dipped gently into the media. The stimulator was set to deliver 5 ms bipolar square pulses of 20mV with 0.5Hz frequency. Muscle contraction and post holder movements were recorded over a period of 5 seconds during stimulation via DinoLite Edge microscope (DinoLite) mounted underneath the posts of each device. Analysis of footage was conducted in Imagej using the MuscleMotion plugin.

#### Immunocytochemistry

After contractility recordings, 3D muscle constructs were fixed in 1% PFA overnight before removal from posts. Constructs were then permeabilised and blocked for 6 hours in TBS 0.05M (1X) pH7.4, 10% FBS, 1% BSA, 0.5% Triton X-100 at 4°C. Prior to overnight incubation with primary antibodies for Titin and MyoH (Table 10) at 4°C in TBS 0.05M (1X) pH7.4, 1% BSA, 0.5% Triton X-100. The following day constructs were washed 6 times in PBS before overnight incubation with secondary antibodies and DAPI at 4°C in TBS 0.05M (1X) pH7.4, 1% BSA, 0.5% Triton X-100. Finally, constructs were mounted on Brand™ Cavity Slides with Fluoromount G (Thermo) prior to imaging.

#### Differentiation and directionality analysis

Images of 3D muscle constructs were captured using an inverted Zeiss confocal with a 40x objective. Z-stacks were taken across the constructs and projected using the SUM function in ImageJ. Single images were isolated, and directionality measured from titin signal using the directionality plugin in ImageJ.

### Rheometric analysis

PDMS 1:10 and 1:20 rheological properties were measured after 60 min of curing at 75°C, by means of the Discovery HR20 rheometer (TA instrument). Samples were analysed at room temperature (25°C), using an 8 mm plate geometry, with a frequency sweep of 1 and an angular frequency from 25.12-0.1 rad/s. Storage (G’) and loss (G’’) moduli were measured for the entire period. Young modulus (E) was calculated following the equation^79^.

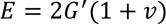

Where 𝐺^′^ is the storage modulus and 𝑣 is the Poisson ratio (0.5).

### Hydrogel culture system

#### Hydrogel composition

Preparation steps started by measuring neuronal culture medium (4 mL) and homogenously mixed with Matrigel (Corning) (1 mL). Hyaluronic acid (5 mg/mL) from Sigma Aldrich was mixed in the above solvent and stirred for overnight (12 hours) at ambient temperature. This was followed by addition of Fibrinogen (45 mg/mL) from Sigma Aldrich and stirred for 5 hours at room temperature. As a final step, Alginate (5% (w/v)) from Sigma Aldrich is added to the above mixture and was allowed to be stirred overnight to obtain a homogenous hydrogel matrix. Motor neuron progenitors were dissociated from a 6-well plate using EDTA in PBS and were gently mixed with the hydrogel matrix (4-5 Million cells/mL) by pipetting them up-down gently for homogenous distribution of cells throughout the hydrogel, this led to the formation of bio-ink. This bio-ink is crosslinked by a 50:50 solution consisting of calcium chloride (CaCl_2_) (1.5% (w/v)) and thrombin (25 U/mL in 0.1% BSA Solution). Bio-ink is allowed to be crosslinked for the duration of 15 minutes at room-temperature. After crosslinking, the bio-ink was washed with PBS solution for 3 times. This was followed by flooding the wells gently through the walls of cell plate with neuronal culture media supplemented with Compound E and kept in incubator at 37 ℃ with 5% CO_2_.

### 2.2 Materials

## 5 Acknowledgments

The Serio lab wishes to acknowledge the support of UK Biotechnology and Biological Sciences Research Council [BB/T011572/1] and the Wellcome Trust [213949/Z/18/Z]. Work in the Tedesco lab is supported by the European Research Council (759108 – HISTOID), AFM-Telethon, BBSRC, Muscular Dystrophy and the NIHR (the views expressed are those of the authors and not necessarily those of the National Health Service, the NIHR, or the Department of Health). Human immortalised myoblasts were provided by the Myoline platform of the Institut de Myologie, Paris, France to the Tedesco lab. This research was funded in whole, or in part, by the Wellcome Trust. For the purpose of Open Access, the author has applied a CC BY public copyright licence to any Author Accepted Manuscript version arising from this submission for the work described here.

## 6. Author Contributions

C.H. and M.Ba. performed and analysed the majority of experiments, from 3D printing establishment and protocol development, cell culture, image acquisition, data analysis and moulds design. C.H., M.Ba., and A.S. wrote the manuscript with feedback from all authors. M.Be. performed Raman spectroscopy experiments and respective data analysis. C.M.G generated a fluorescent motor neuron line used in this project. P.S, K.O’T. contributed to astrocyte and cortical cell culture for the multicellular experiments. V.L. performed culture, stimulation and staining of 3D muscle constructs and collected the images with support from N.K.. N.K. and E.G. contributed to the optimisation of 3D constructs design for muscle tissue engineering. G.K. performed SEM imaging. F.S.T. and A.I. provided feedback on data analysis and manuscript. Resources were provided by A.S., A.I, F.S.T. Funding for this project was provided by A.S.

## 7. Supplementary Information

**Figure S1:**
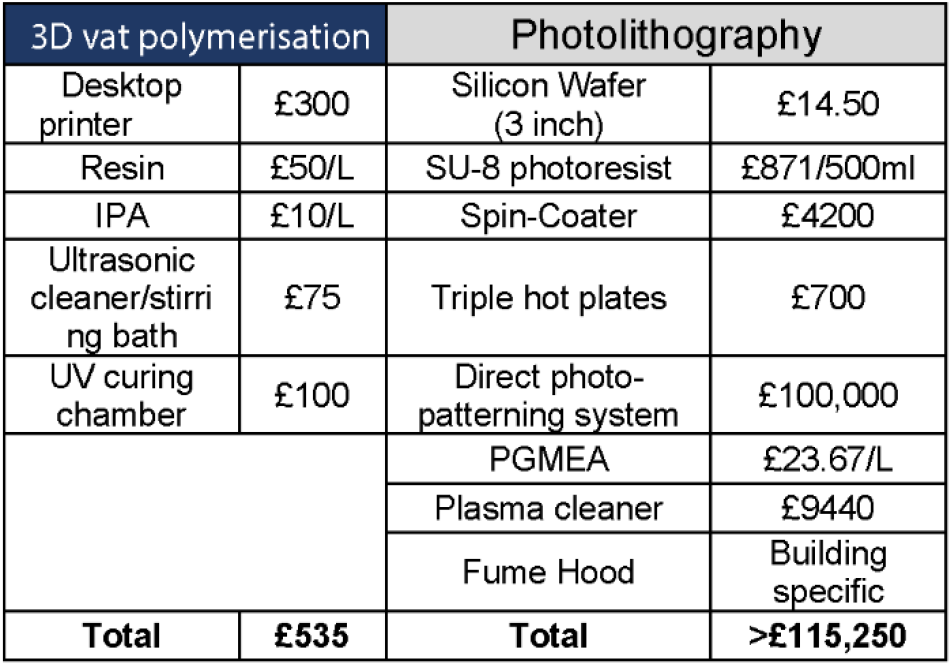
Comparison of costs for 3D vat polymerization printing and photolithography with all the required equipment and materials but excluding personnel training costs.

**Supplementary Diagram 1:**
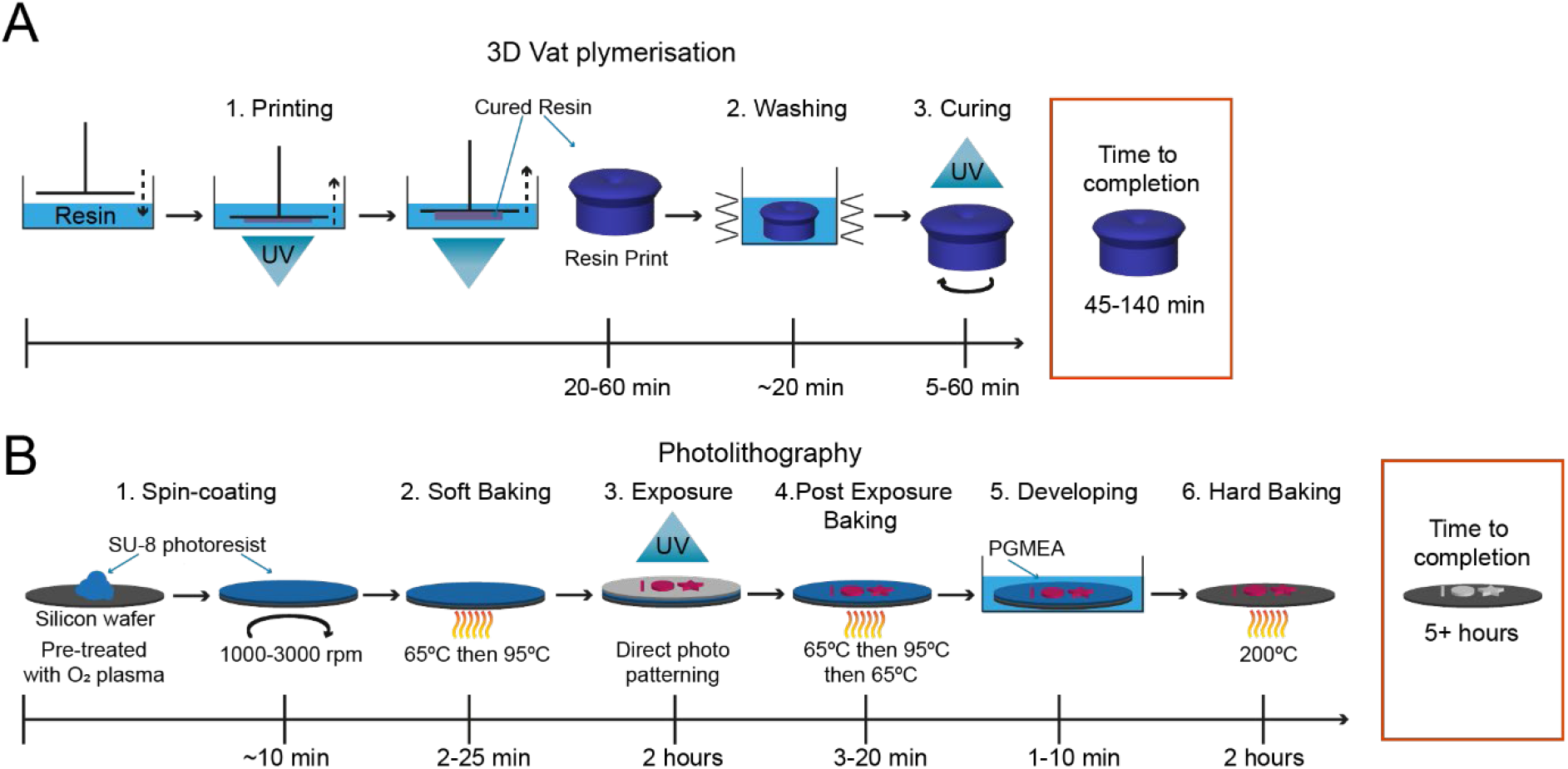
Description of 3D vat polymerization and soft lithography with estimated durations. (A) Schematic overview of the UV resin vat polymerisation process and post-processing with a time estimate for production. Printing times range depend on the printed volume but for cell culture device range from 10min to 1h. (B) Schematic overview of photolithography and an estimated time to completion of a print. Print time is a estimate for feature creation across the whole silicon wafer and alters in between prints, depending on the design. Overall, UV resin vat polymerisation is faster compared to photolithography, as it uses whole field illumination and not single point illumination as photolithography.

**Figure S2:**
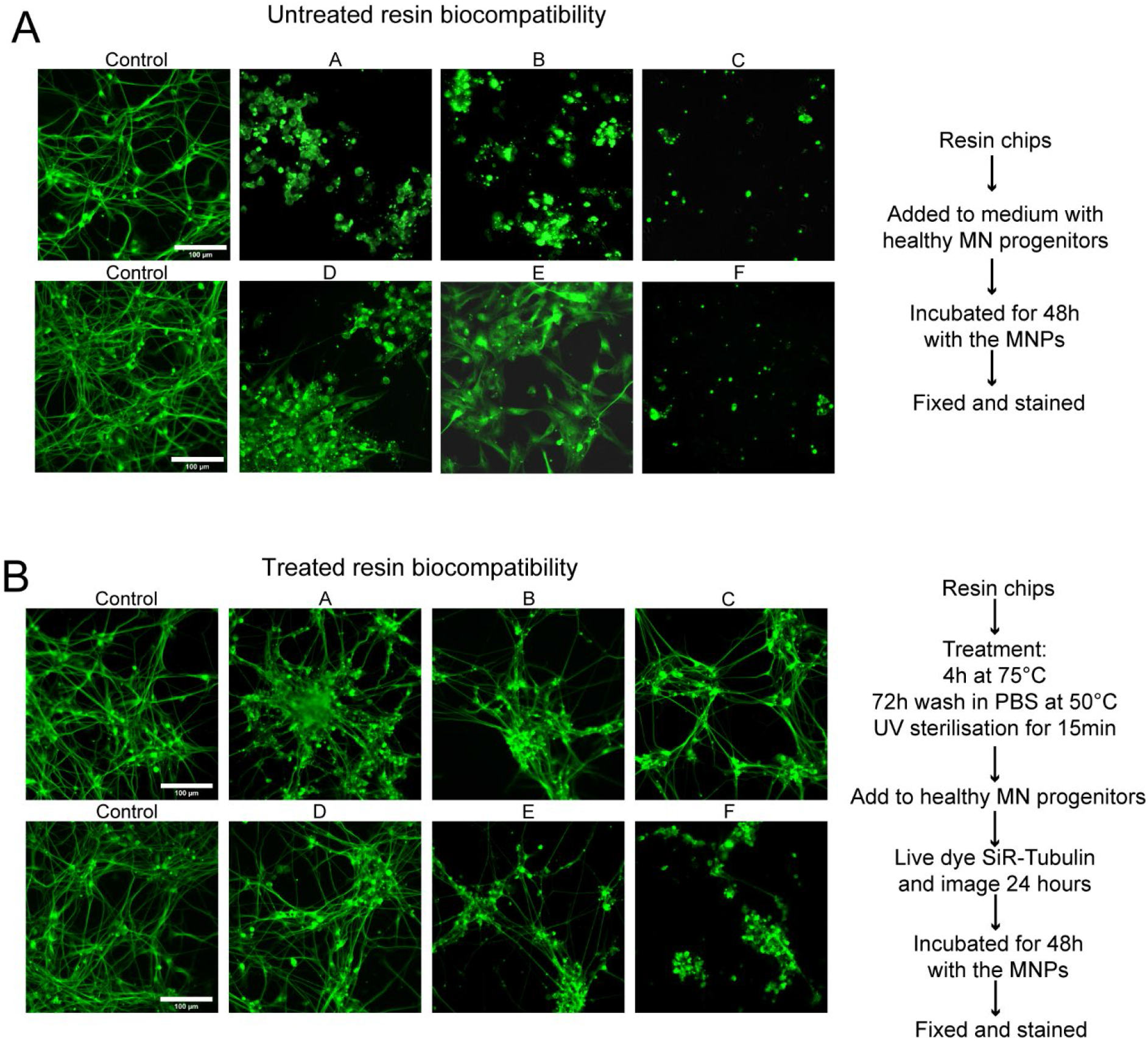
SLA resins themselves are toxic with and without pre-treatment. (A) Chips of 6 resins were added to cultures of motor neuron progenitors pre-stained with SiR – tubulin live dye, incubated for 48 hours before imaging. Representative images of SiR live dye-stained motor neurons after 48 hours in culture with resins compared to 2 control wells. (B) Chips of 6 resins were treated with extra processing steps to improve biocompatibility, bake 4 hours at 75°C, wash in PBS 72 hours at 50°C, UV sterilise 15 mins before being added to cultures of motor neuron progenitors pre-stained with SiR-tubulin cytoskeletal live dye, incubated for 48hrs before imaging. Representative images of SiR live dye-stained motor neurons after 48 hours in culture with treated resins compared to 2 control wells. (C) Representative time lapse videos of SiR live dye-stained motor neurons during the first 24 hours in culture with treated resin chips compared to 2 control wells.

**Figure S3:**
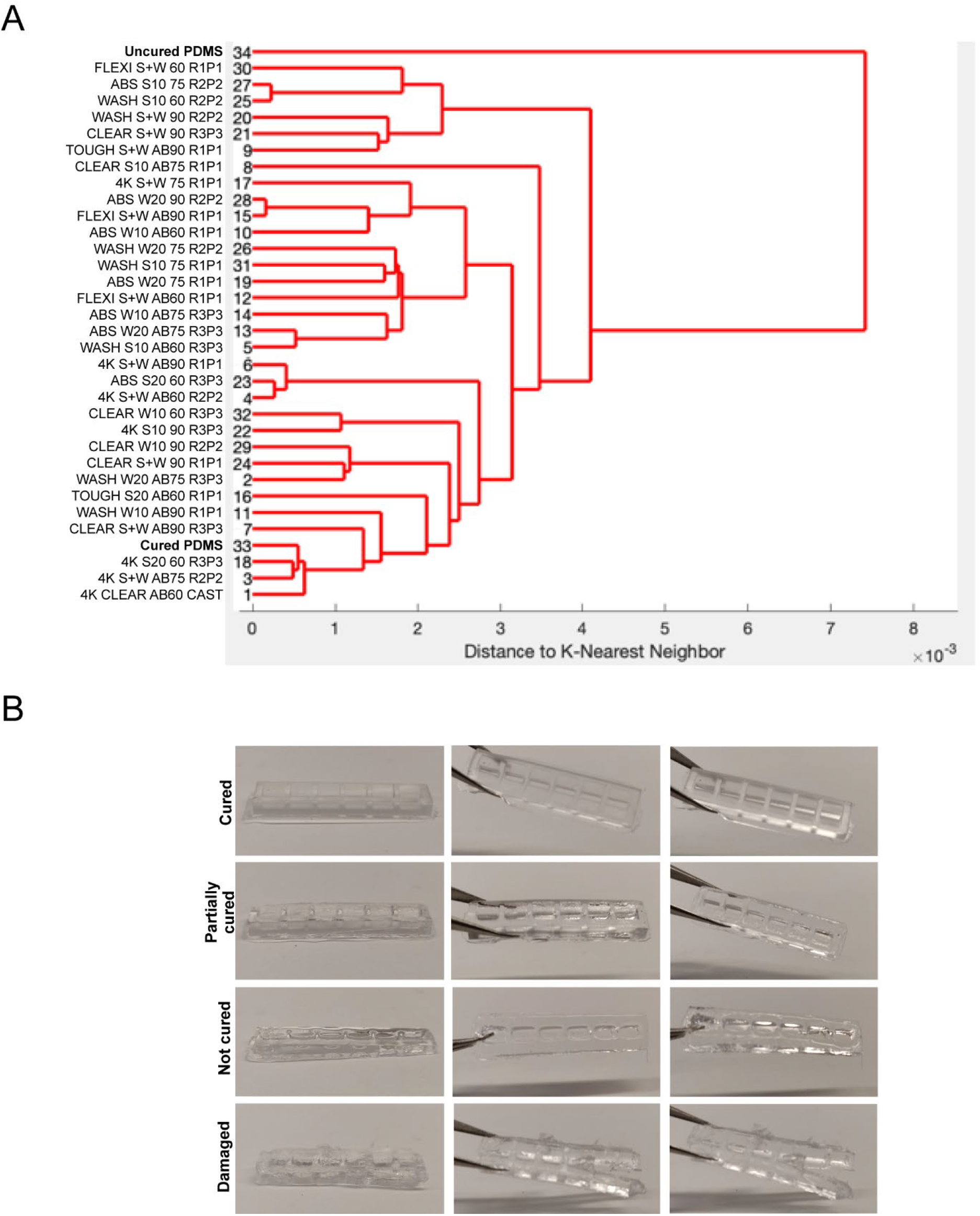
Quantitation of PDMS curing on 3D printed moulds. (A) Dendrogram of the spectral similarity of PDMS casts from 3D printed moulds fabricated with 6 commercially available resins, washed with 5 conditions (S+W = sonicate 10 mins, wash 10 mins, S10 = Sonicate 10 mins, S20 = sonicate 20 mins, W10 = Wash 10 mins, W20 = Wash 20 mins), either untreated or coated with airbrush (AB) and cured at 3 different temperatures (60), (75), (90), compared to samples of uncured and cured PDMS. Replicate and print number (RxPx) for each condition. (B) Heterogeneity of PDMS cast curing from 3D printed moulds by observation ranging from ‘cured’, ‘partially cured’, ‘not cured’, ‘damaged’.

**Figure S4:**
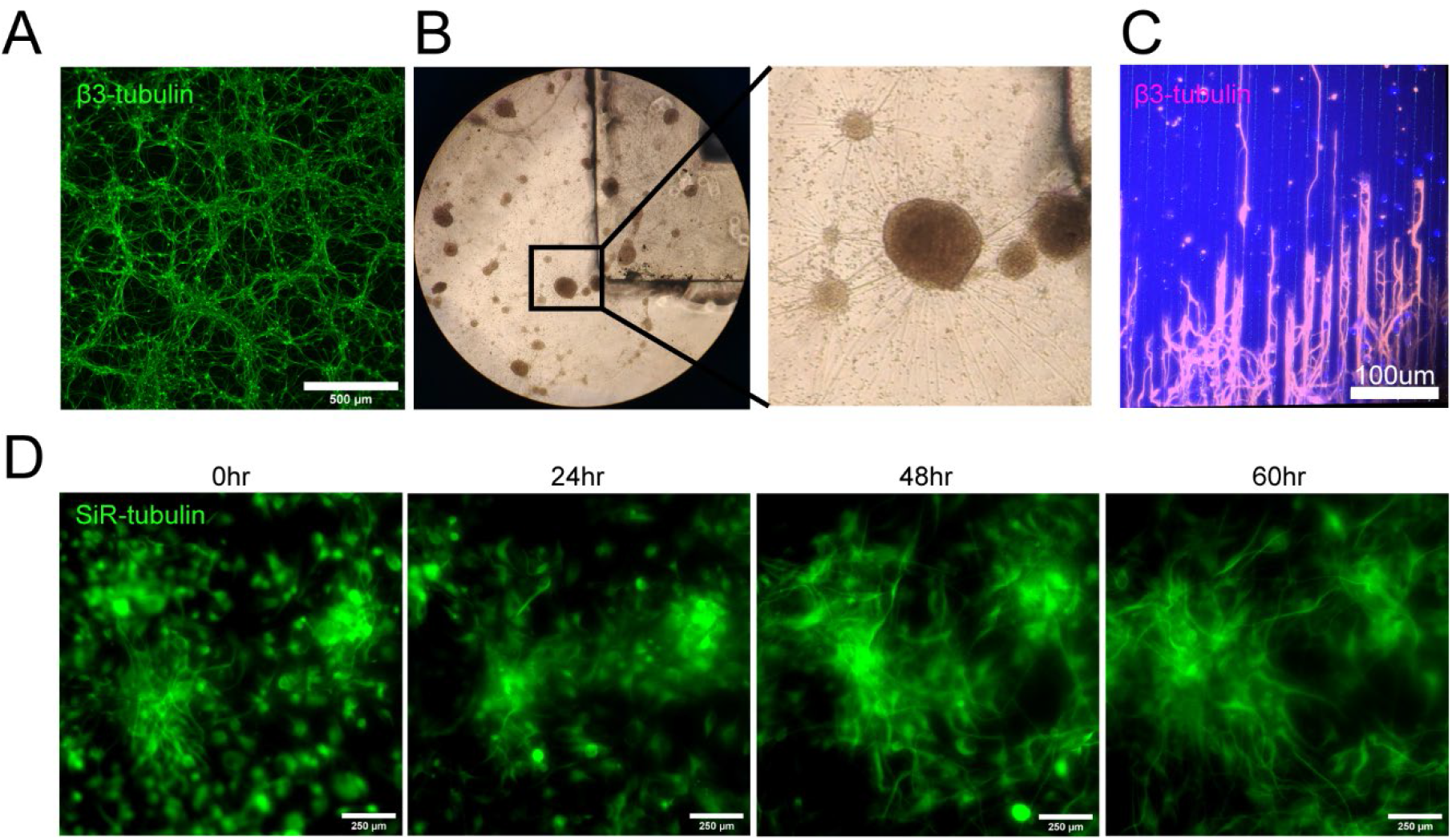
Biocompatibility of PDMS cast in 3D printed moulds with cells (timelapse and long axons) (A) Representative β-III Tubulin stained differentiated motor neurons cultured on a flat substrate. (B) Brightfield images of differentiated motor neurons with a PDMS cast from a 3D printed mould in the culture medium. (C) β-III Tubulin stained motor neuron long axons seeded on PDMS microfabricated substrate. (D) Snapshots from 60-hour time-lapse of SiR – tubulin live dye stained motor neuron differentiation on PDMS substrate cast in a 3D printed mould.

**Figure S5:**
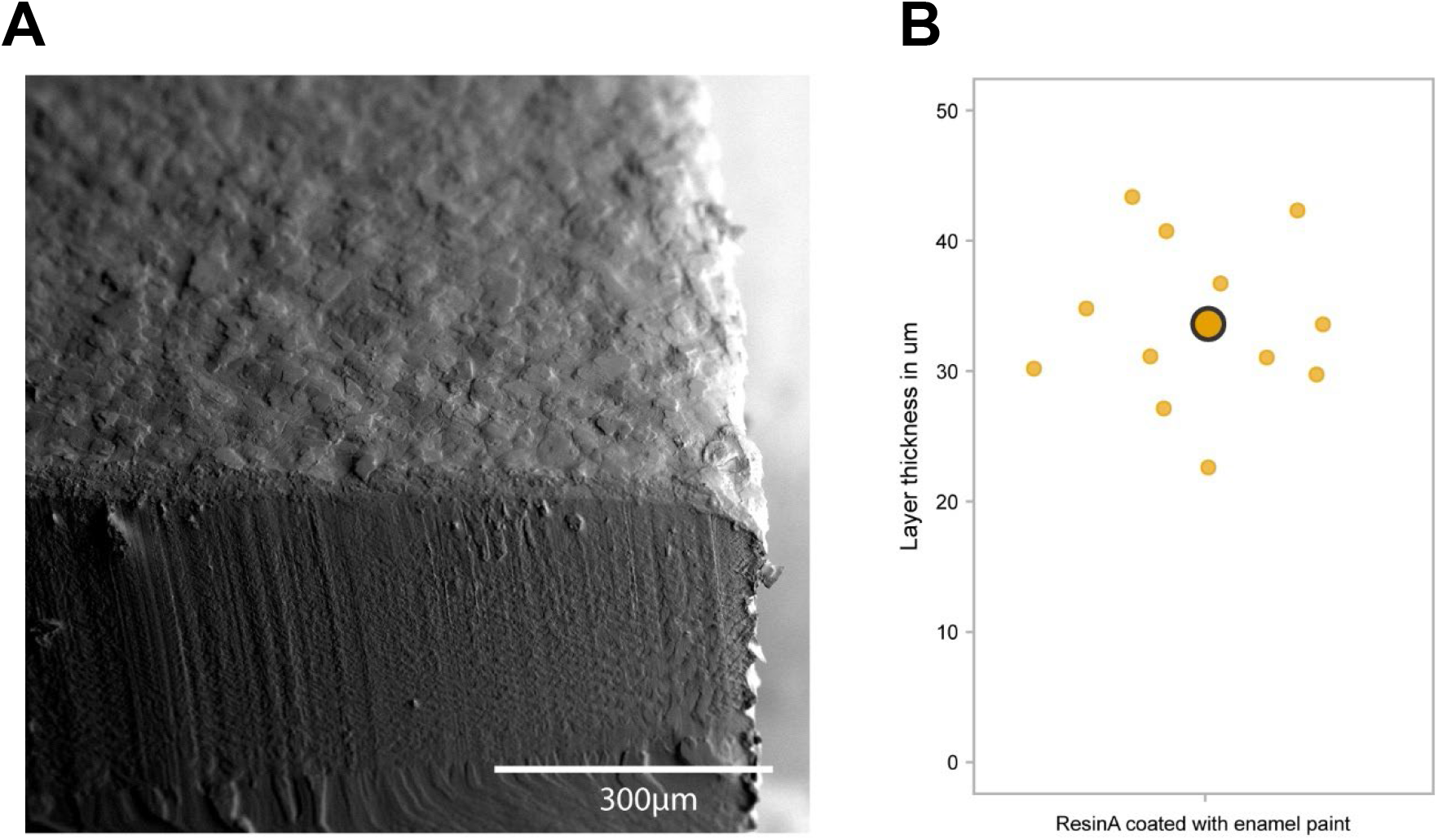
Analysis of paint layer thickness. (A) Representative SEM image of a resin A print with airbrushed enamel paint (B) Analysis of paint layer thickness on 3D vat printed moulds

**Figure S6:**
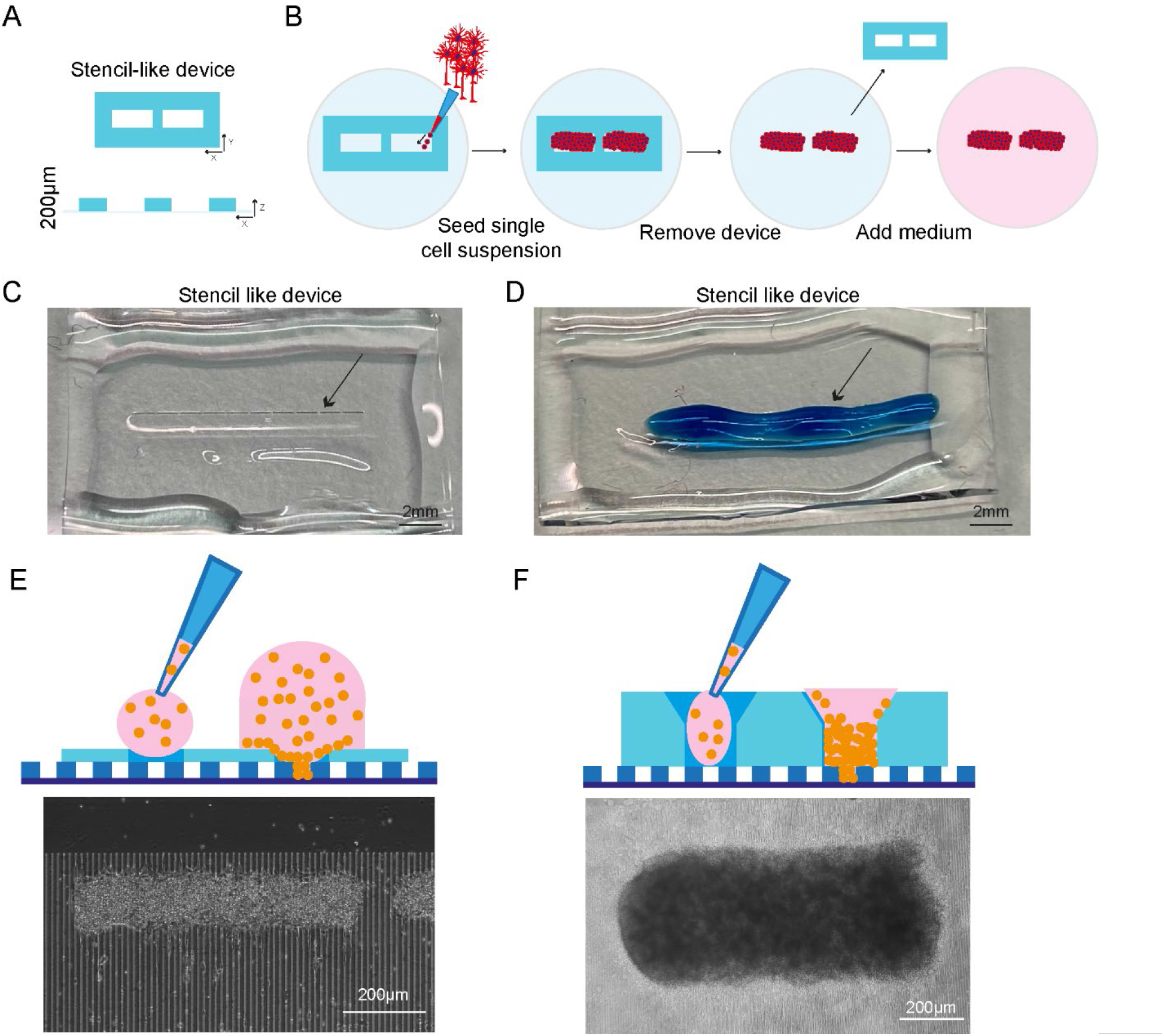
PDMS stencil devices manufactured using photolithography. (A) Dimensions of the stencil-like device with photolithography (B) Schematic overview of the seeding strategy with stencil-like devices (C) Representative image of a stencil-like device with pockets for cell seeding (arrow) (D) Demonstration of cell seeding using food coloring as a ‘single-cell suspension’. The liquid was manually pipetted on the pockets of the device (arrow). Limited volume can be used. (E) Representative brightfield image of single motoneurons seeded using a stencil device manufactured using photolithography and a schematic showing the seeding process as well as the position of the cells. (F) Representative brightfield image of single motoneurons seeded using a plating device manufactured using SOL3D and a schematic showing the seeding process as well as the position of the cells.

**Figure S7:**
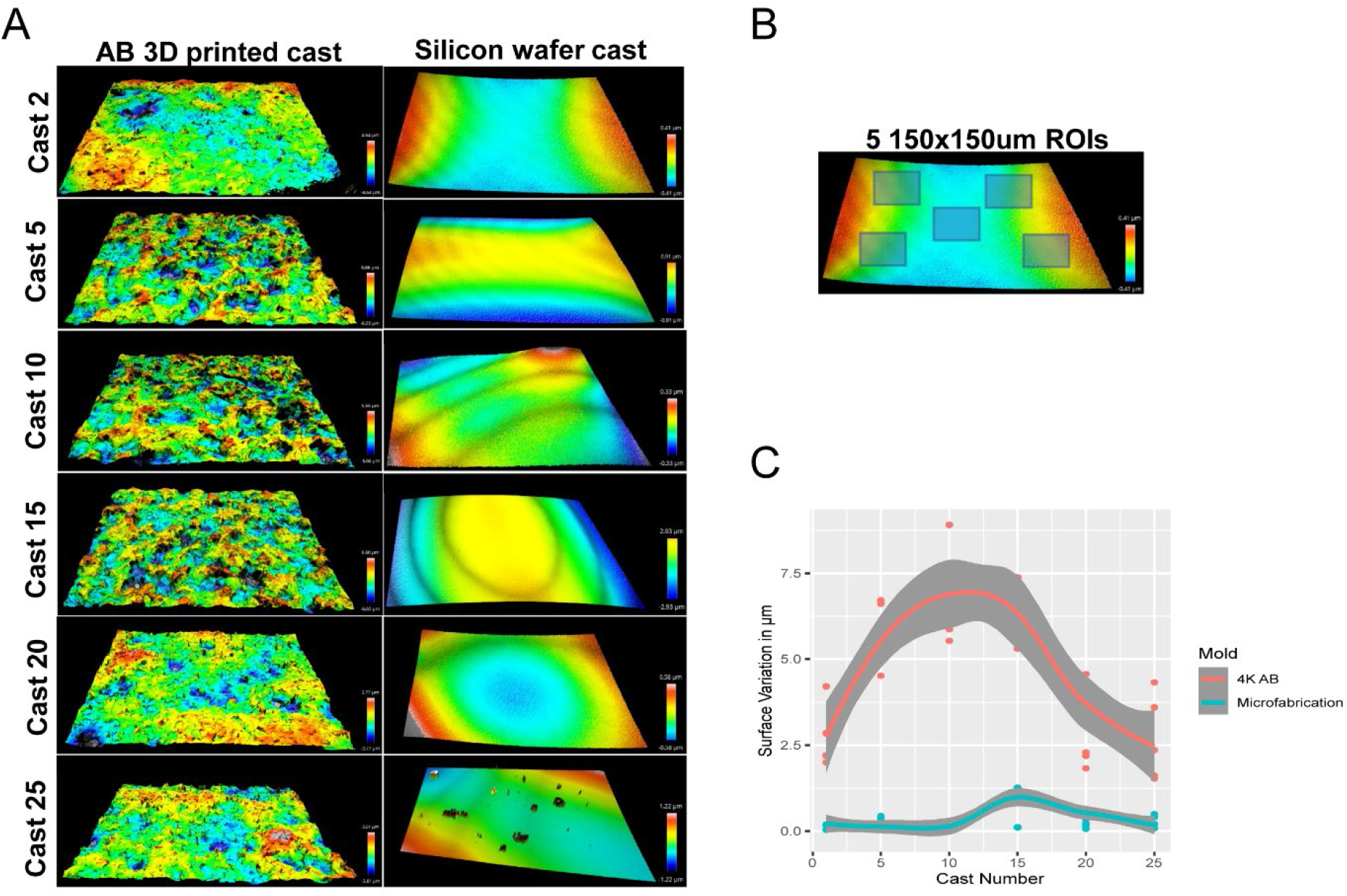
The surface of 3D prints is rough. (A) Representative optical profiles from PDMS casts demoulded from a single airbrushed 3D printed mould and a single microfabricated mould. (B) Representation of 5 ROI selection for quantification of surface roughness on PDMS casts. (C) Quantitation of surface roughness of PDMS casts from the same device over time (25 simultaneous casts) between an airbrushed 3D printed mould and a microfabricated mould.

**Figure S8:**
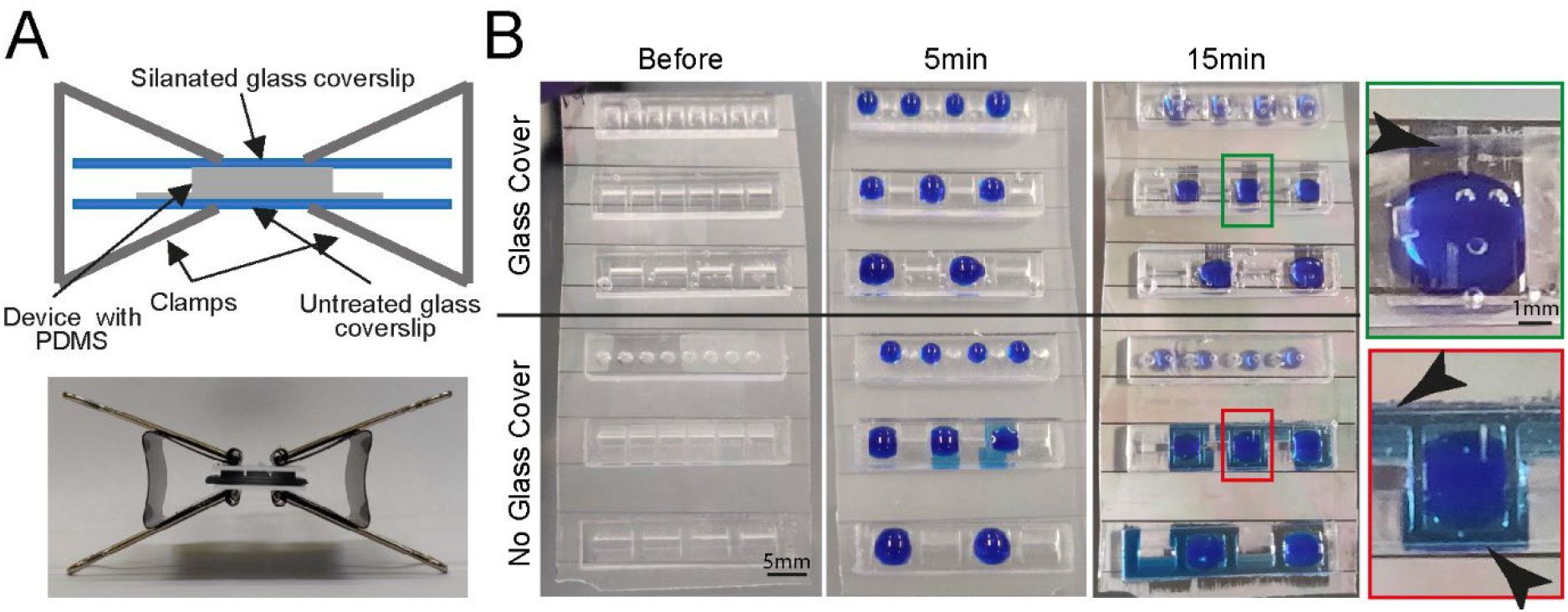
Fluidic seal for cell plating devices. (A) Schematic (top) and representative image (bottom) of clamping approach to ensure fluid seal when devices are placed on PDMS microgroove substrates. (B) Comparison of liquid seal integrity of clamping strategy (top) compared to open curing (bottom) on PDMS microgroove substrate with different well sizes and shapes using dyed liquid. Successful sealing of devices cast with a glass cover (**Green zoom)**. Dye spreads throughout device and grooves using open cured (**Red zoom)**. Arrows highlight liquid spreading.

**Figure S9:**
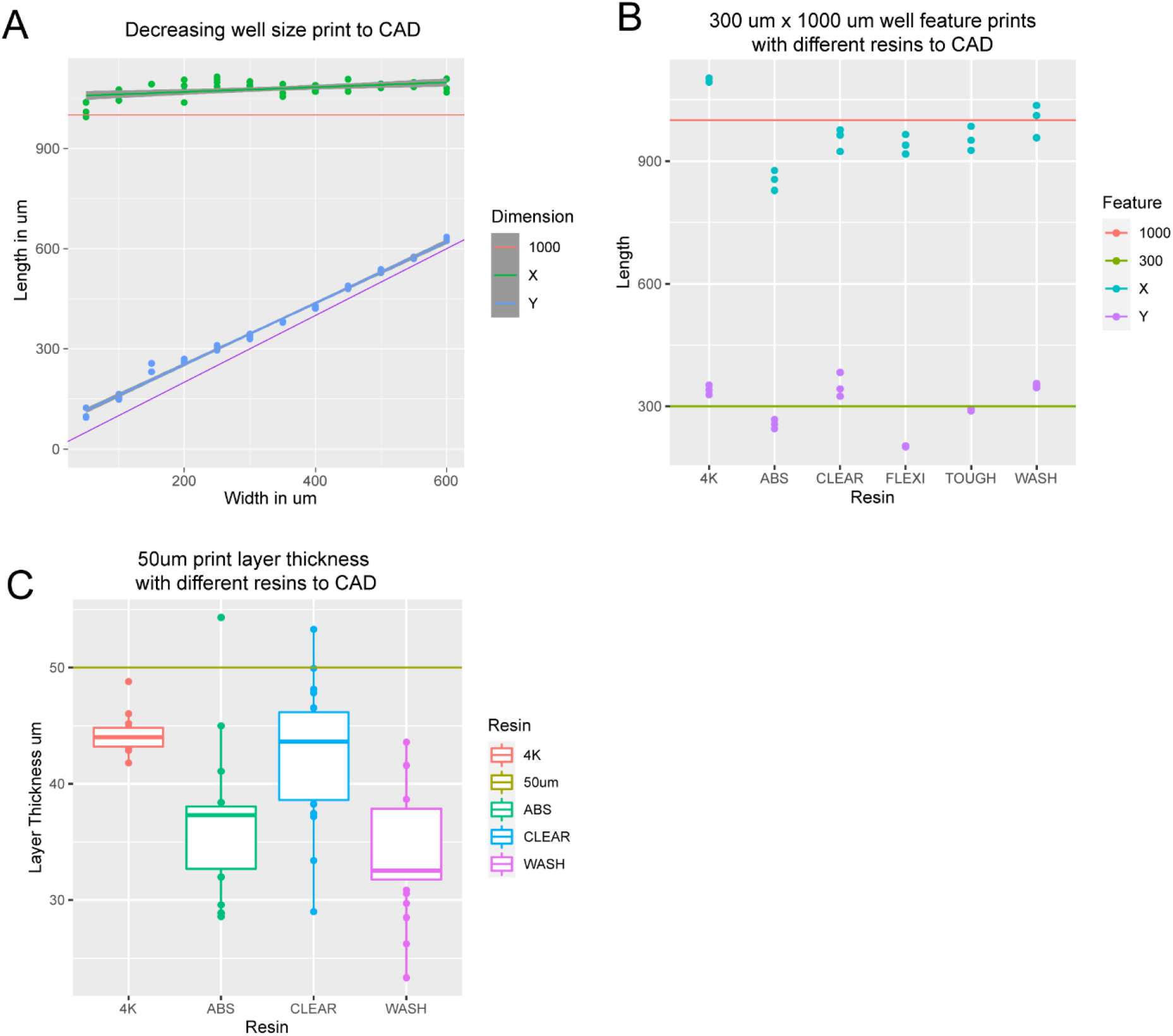
3D print dimensions are homogenous. (A) Graph comparing X and Y dimensions of 3D printed constructs to CAD specifications in a single device with well dimensions ranging from 600 µm x 1000 µm to 50 µm x 1000 µm. (B) Graph comparing X and Y dimensions of 300 µm x 1000 µm features on 3D printed constructs to CAD specifications for 6 commercially available resins printed on 2 3D printers at manufacturer default settings with a 50 µm layer thickness. (C) Graph comparing the actual layer thickness of 3D printed constructs to CAD specifications for 6 commercially available resins printed on 2 SLA 3D printers at manufacturer default settings with a 50 µm layer thickness.

**Figure S10:**
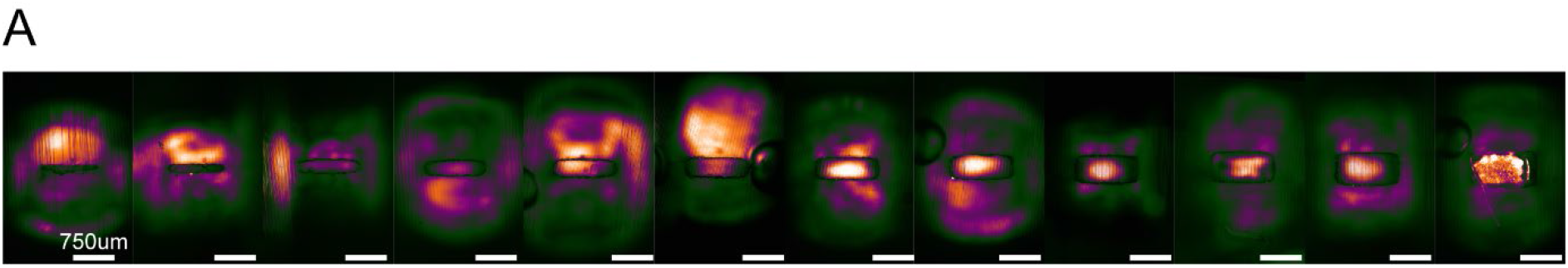
Non-Plasma devices cannot be used for seeding in microwells. (A) Representative SiR-Tubulin images of motor neuron progenitors seeded in microwells ranging from 600 µm x 1000 µm to 50 µm x 1000 µm. Difference in colour indicates depth in focal plane where cells do not reach the micropatterned substrate below.

**Figure S11:**
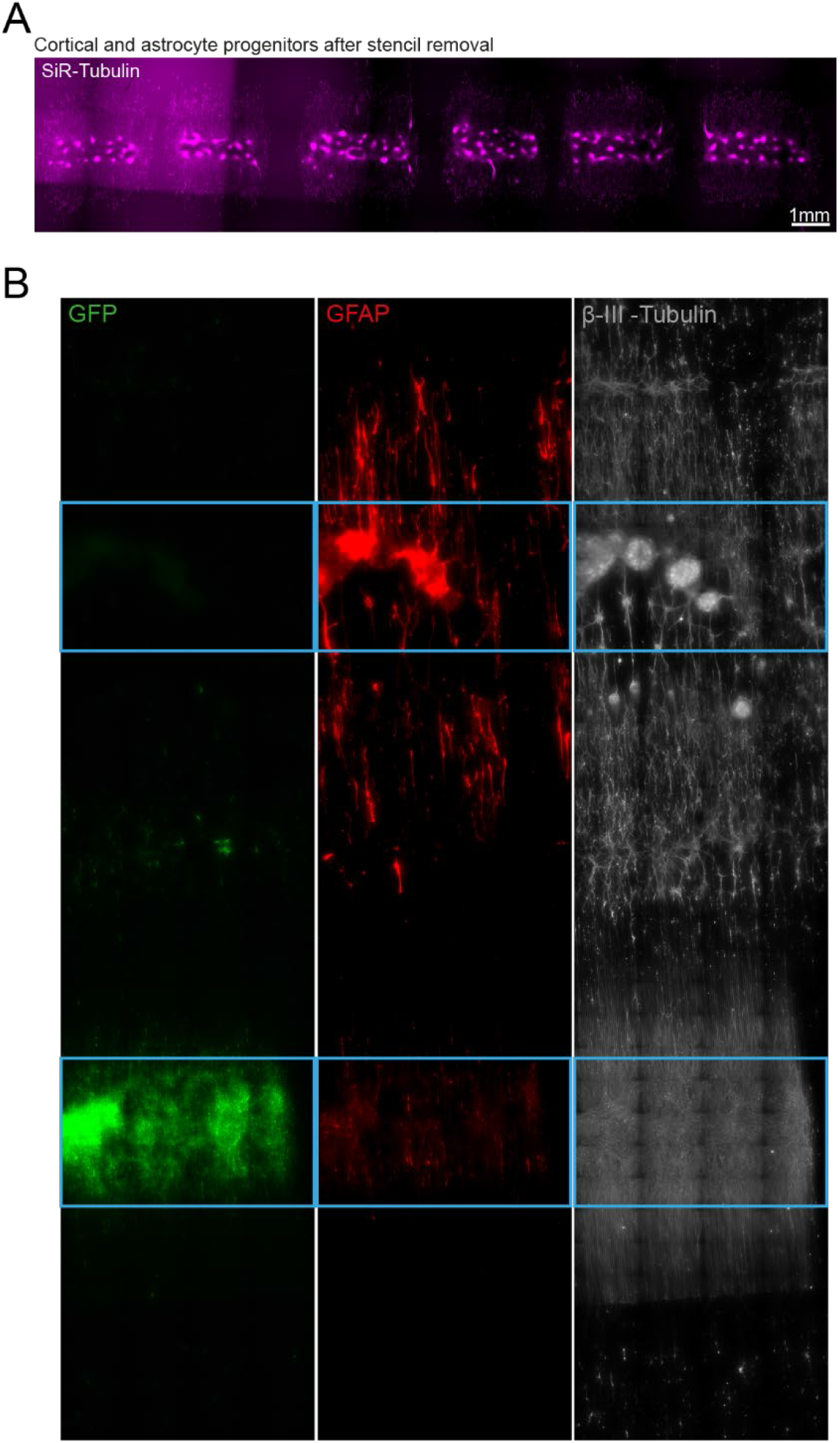
Spatio-temporal control over cell seeding in an open well. (A) Representative SiR tubulin live cell dye fluorescence images of astrocyte and cortical progenitors. Images were captured 2 days following device removal (B) Channel split of complete circuit after 19 days of culture. GFP transfected motor neurons = green, Glial Fibrillary Acidic Protein (GFAP) identifies astrocytes, β-III – Tubulin identifies cortical neurons and the tubulin in GFP+ motor neurons. Blue device well shapes overlaid for illustrative purposes.

**Figure S12:**
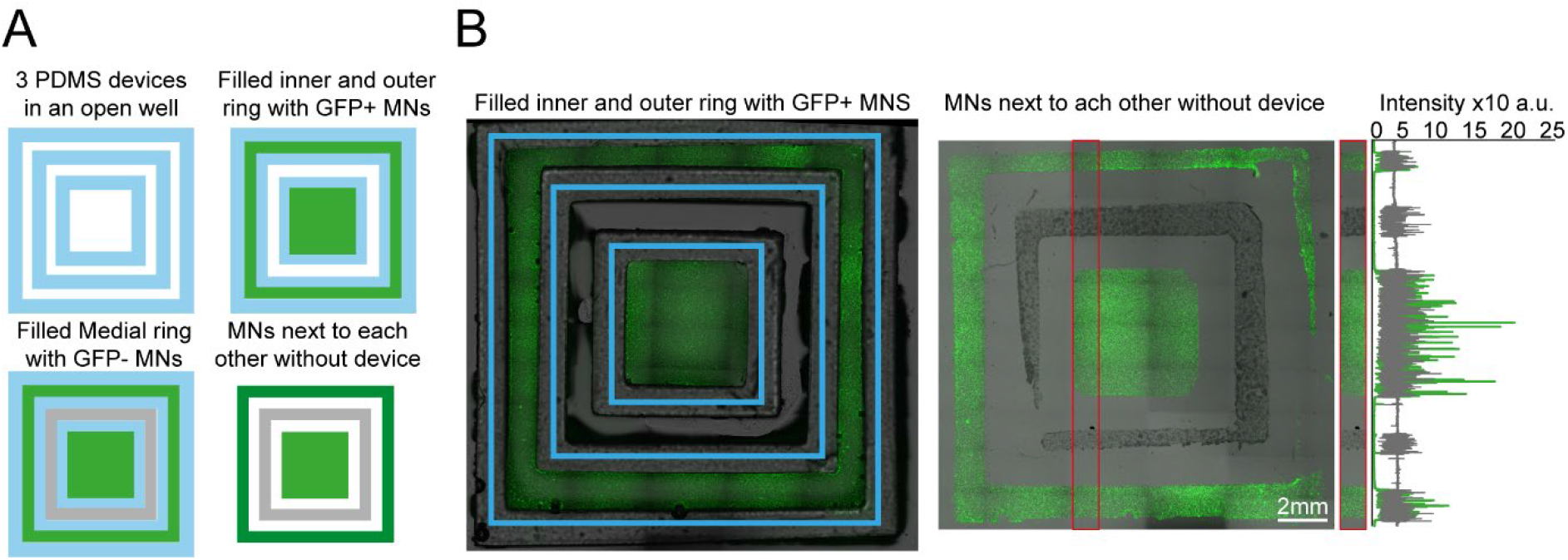
Plating devices enable manual segregated seeding of different cell types in the same well, device or multiple devices. (A) Multiple PDMS casts from 3D printed devices can be seeded in the same well and seeded with different cell types at different time points. Schematic overview of the multi-device protocol for seeding GFP and non GFP+ motor neurons at different time points in 3 devices in the same well. (B) Representative fluorescence images of cells GFP+ motor neurons seeded in the inner and outer rings of the 3 devices and imaged with devices still on (left) and after seeding of the second non-GFP+ motor neurons in the medial ring after device stripping (right), Representative line profile of imaged cell fluorescence showing segregation of individual populations to their designated rings

**Figure S13:**
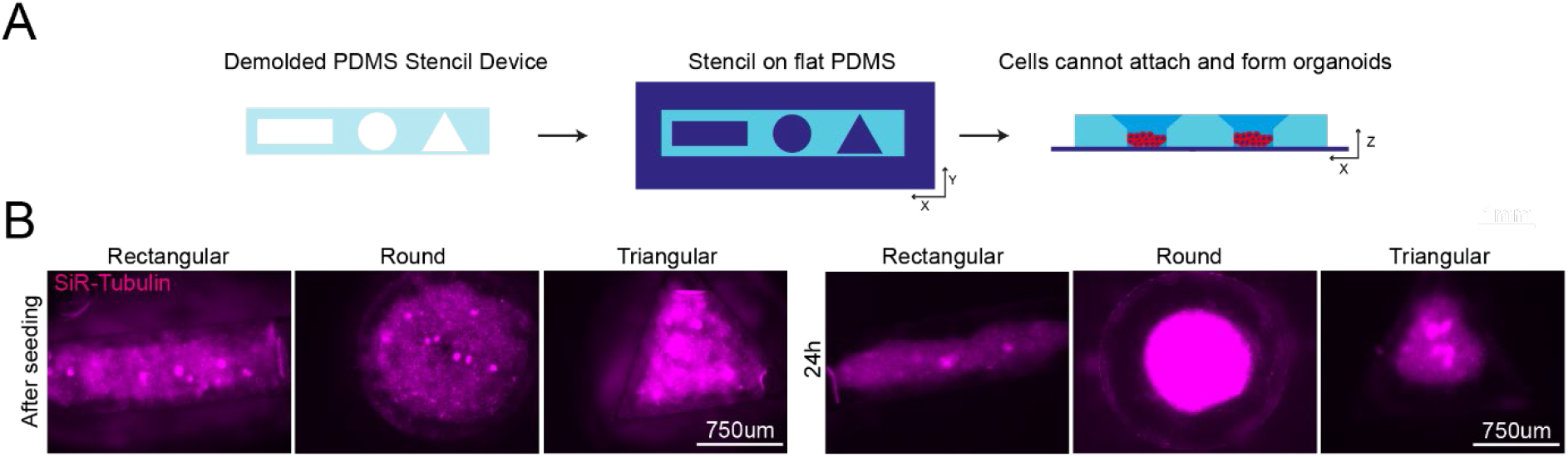
Plating devices enable geometric manipulation of aggregoid cultures in 2D and 3D. (A) Schematic overview of protocol for manipulating aggregate geometry in combination with existing microgroove and flat substrates. (B) Representative images of SiR-tubulin live-cell-stained motor neuron aggregoids at day 2 (top) and β-III Tubulin staining after 11 days (bottom) of culture. (C) Boxplot of aggregoid aspect ratio fold change by shape between CAD (blue line) and day 2 of culture from β-III-Tubulin channel (top) (D) Boxplot of aggregoid area fold change by shape between CAD (blue line) and day 2 of culture from β-III Tubulin channel. (E) Representative images of SiR-tubulin stained cortical aggregates after seeding and 24h later in different geometry stencil devices, round (left), triangular (middle) and rectangular (right).

**Figure S14:**
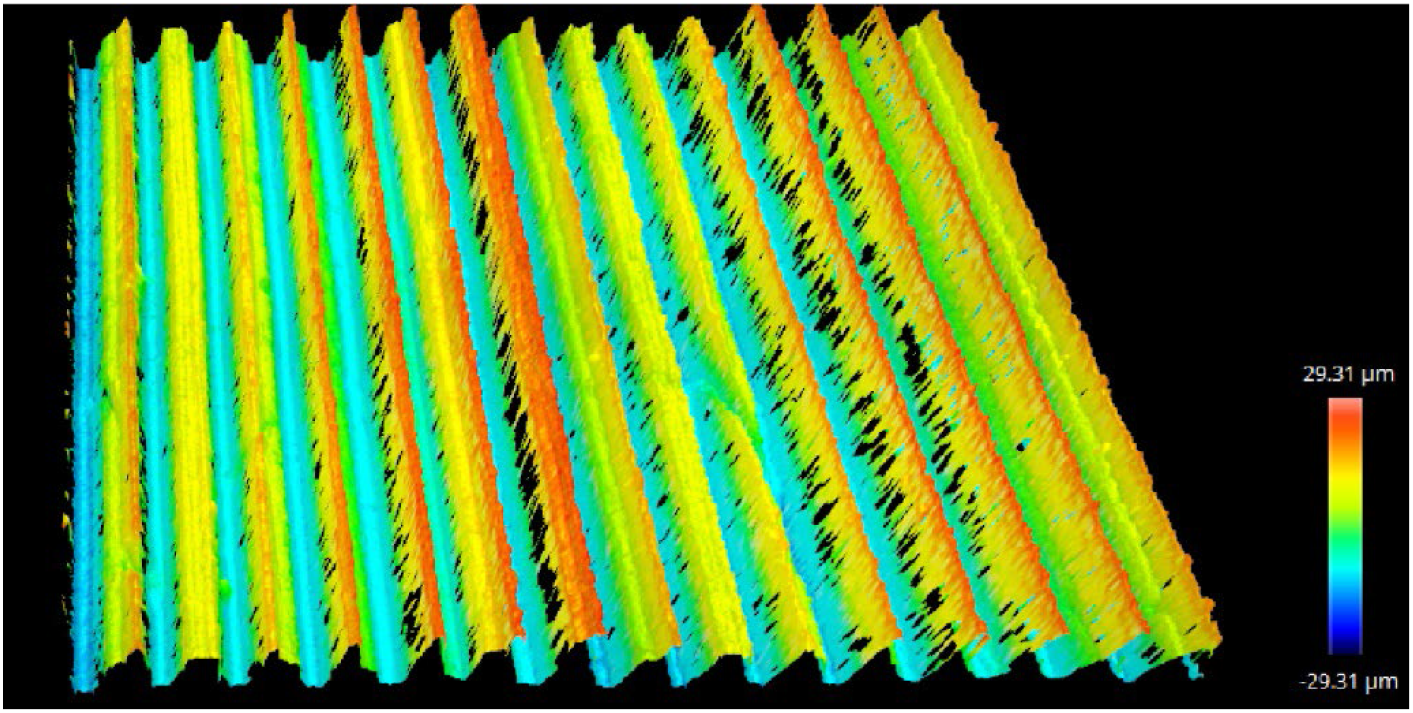
Representative image of an optical profile from SOLID manufactured triangular grooves.

**Figure S15:**
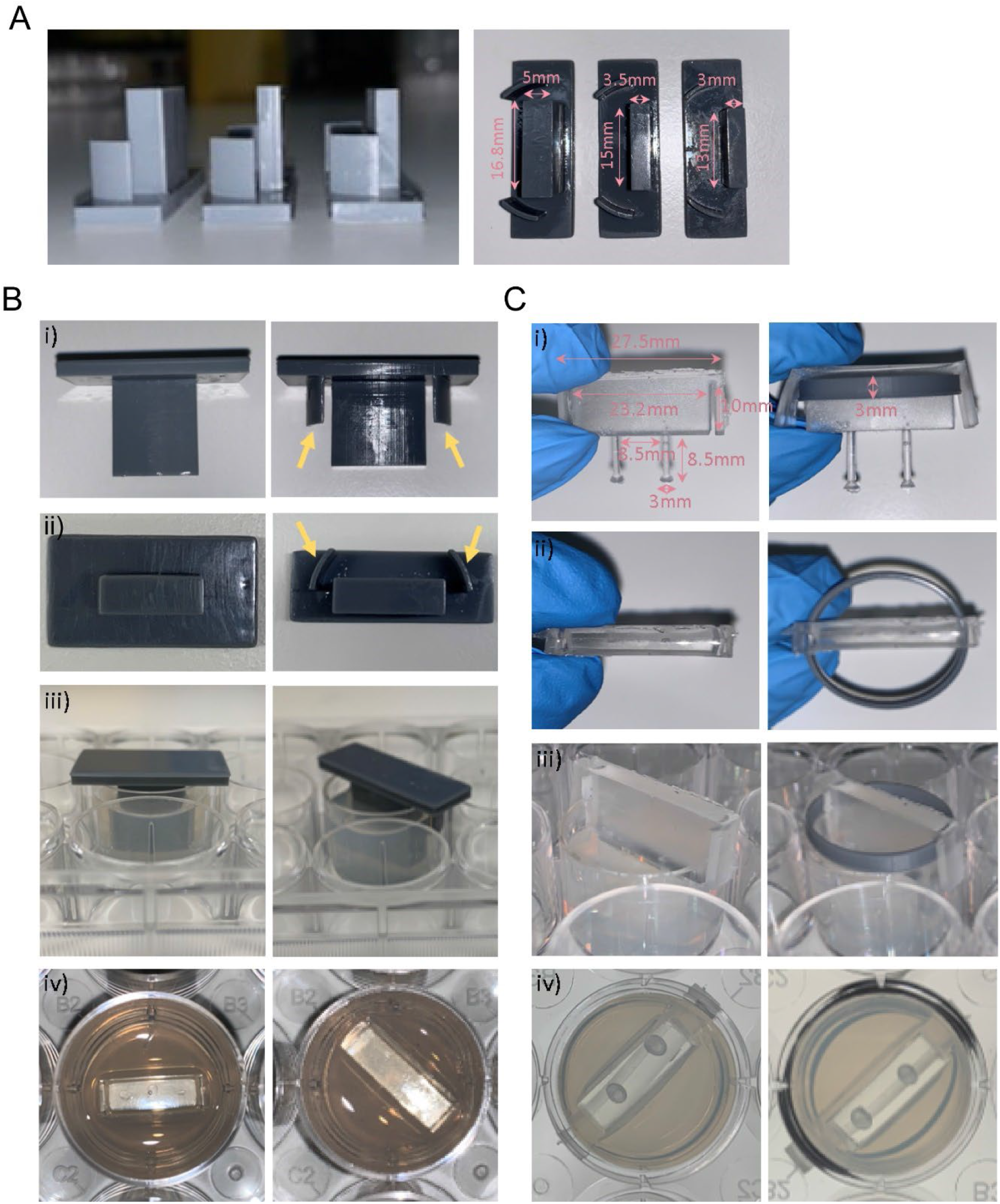
Optimisation and protocol for 3D muscle culture using PDMS constructs from 3D printed moulds. (A) Images show three different mould sizes tested to optimise the volume of the hydrogel mix at a side and bottom view. (Bi-iii) Images showing moulds with (yellow arrows) and without grooves at various perspectives. (Biv) Image shows top view of agarose once mould is removed. (Ci-ii) Images show posts with and without a ring placed underneath the arms from a front and top view. Rings were designed to match the size of one well of a 12-well plate. (Ciii) Images showing posts inserted into agarose moulds at a side view and bottom view from underneath the plate, rings allow posts to be inserted at a specific height.

**Figure S16:**
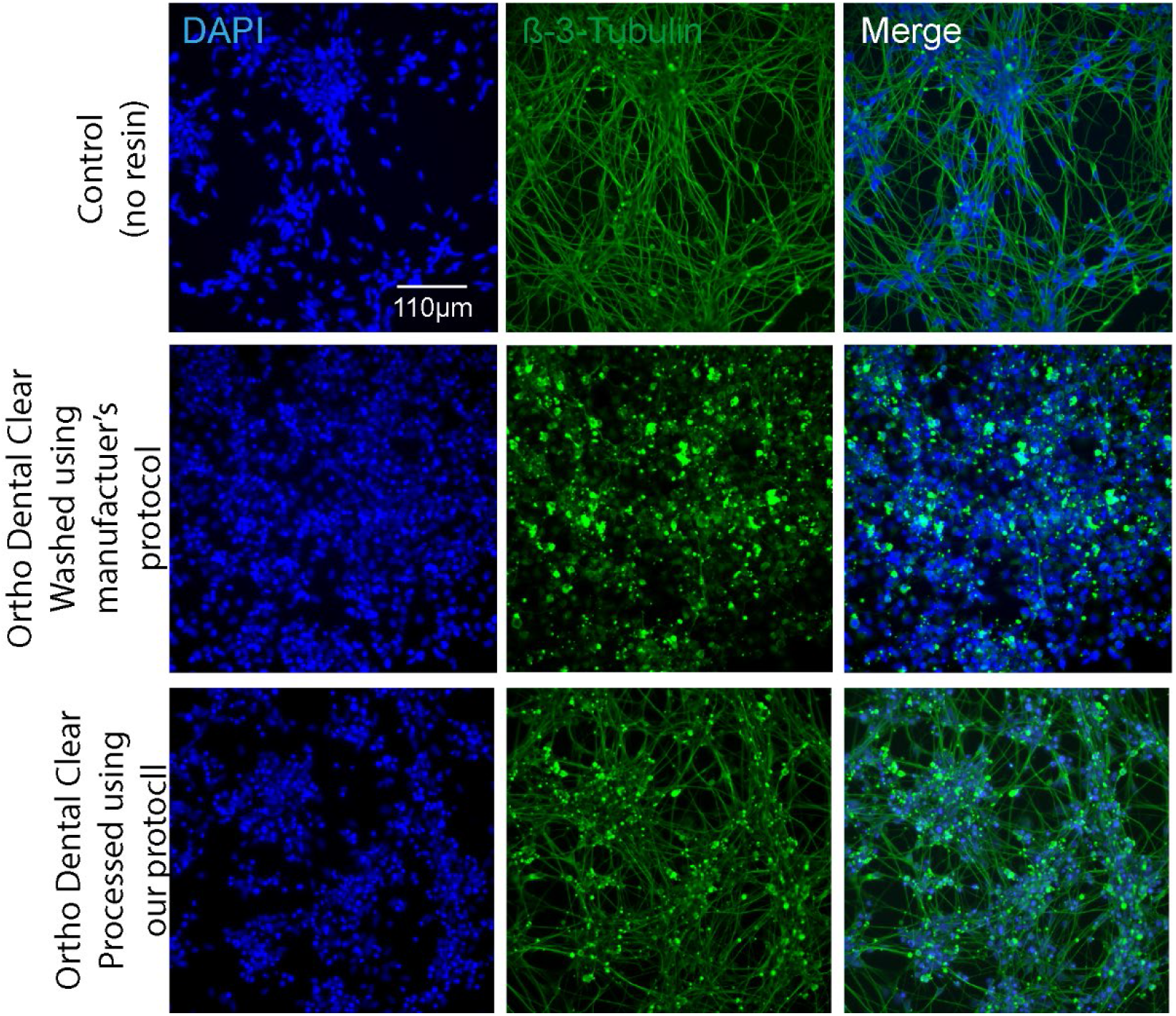
Effects of co-culture with a biocompatible resin on iPSC-derived MNs. iPSC-derived MNs were plated at equal densities. Then a 3D-printed well-sized cylinder of ortho-clear resin was added to the well and incubated for four days to identify the toxic effects of the biocompatible resin. The resin was either pretreated with the recommended manufacturer’s protocol or our pipeline. Cells were stained for β-3-tubulin and DAPI.

**Figure S17:**
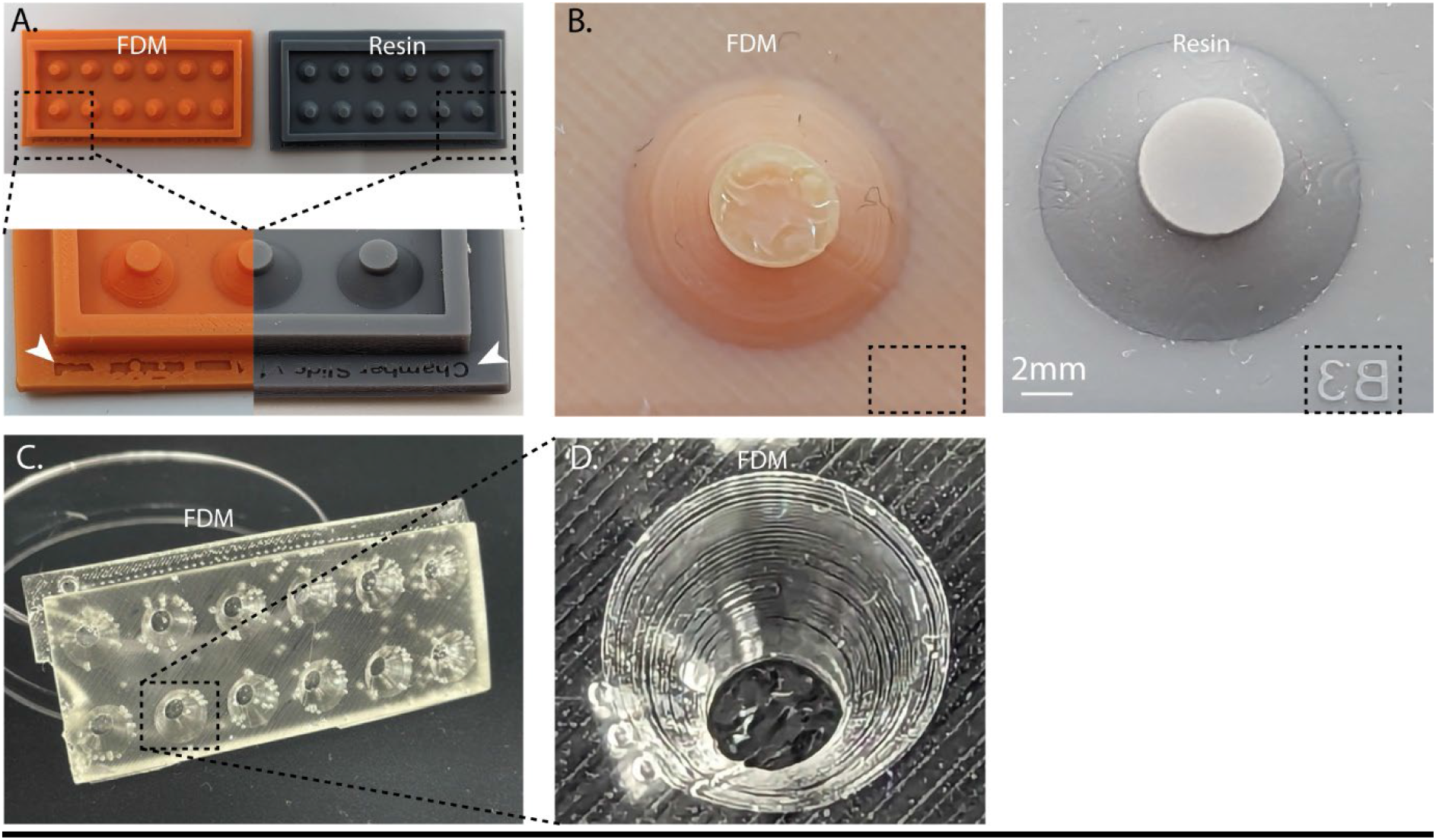
Comparison of feature quality between FDM and Vat polymerized samples. (A) Chamber devices used in Figure 5 were manufactured using a FDM (Bambu Labs, 0,4mm nozzle with 0.16mm settings) printer and a 3D UV-resin printer (Phrozen mini 4k, 50um settings). Zoom in images show a clear constructed writing for the UV resin printer, lacking for the FDM sample. (B) Representative images highlight the flat surface of resin printed samples, especially on the flat top of the pillar. The dotted box highlights the absence of the well labelling in FMD prints, and a clear label in the resin sample. (C-D) PDMS cast of the FDM mould, clearly highlights the large print and build lines of the sample.

